# Robust and replicable effects of ageing on resting state brain electrophysiology measured with MEG

**DOI:** 10.1101/2025.08.01.668093

**Authors:** Andrew J. Quinn, Jemma Pitt, Oliver Kohl, Chetan Gohil, Mats W.J. van Es, Anna C. Nobre, Mark W. Woolrich

## Abstract

Non-invasive recordings using brain electrophysiology provide credible insights into decline in neuronal functioning with age. New approaches are required to translate these results into compelling clinical metrics and meet the global challenge of preserving brain health in ageing. Changes in neuronal dynamics with ageing are observable in the power spectra of EEG and MEG recordings. Highly promising candidates for electrophysiological markers have been identified, but progress is hindered by substantial methodological variability across studies. This makes it challenging to establish a clear consensus on the frequency, location, and direction of any single reported effect within the larger body of research. We estimate a full-frequency whole-head profile of the ageing effect on eyes-closed resting-state MEG using the GLM-Spectrum. This data representation is easily sharable, facilitates meta-analyses and provides a framework for estimating statistical power of age effects for future study planning. We use this to show that the effect of age replicates across open-access MEG datasets and is robust to modelling of common covariates. Distinct components within the full frequency profile have different effect sizes, indicating that sample-size planning for ageing effects must consider the specific features of interest. The frequency profile of ageing is strongly robust to a range of common covariates and partially robust to modelling of grey matter volume. We establish that what seems a well-powered study may become underpowered when analyses target an age effect that is linearly separable from an age-relevant covariate such as grey matter volume. These results provide a pathway towards formal comparison and assessment of candidate markers for brain health in ageing.

**Graphical Abstract:** 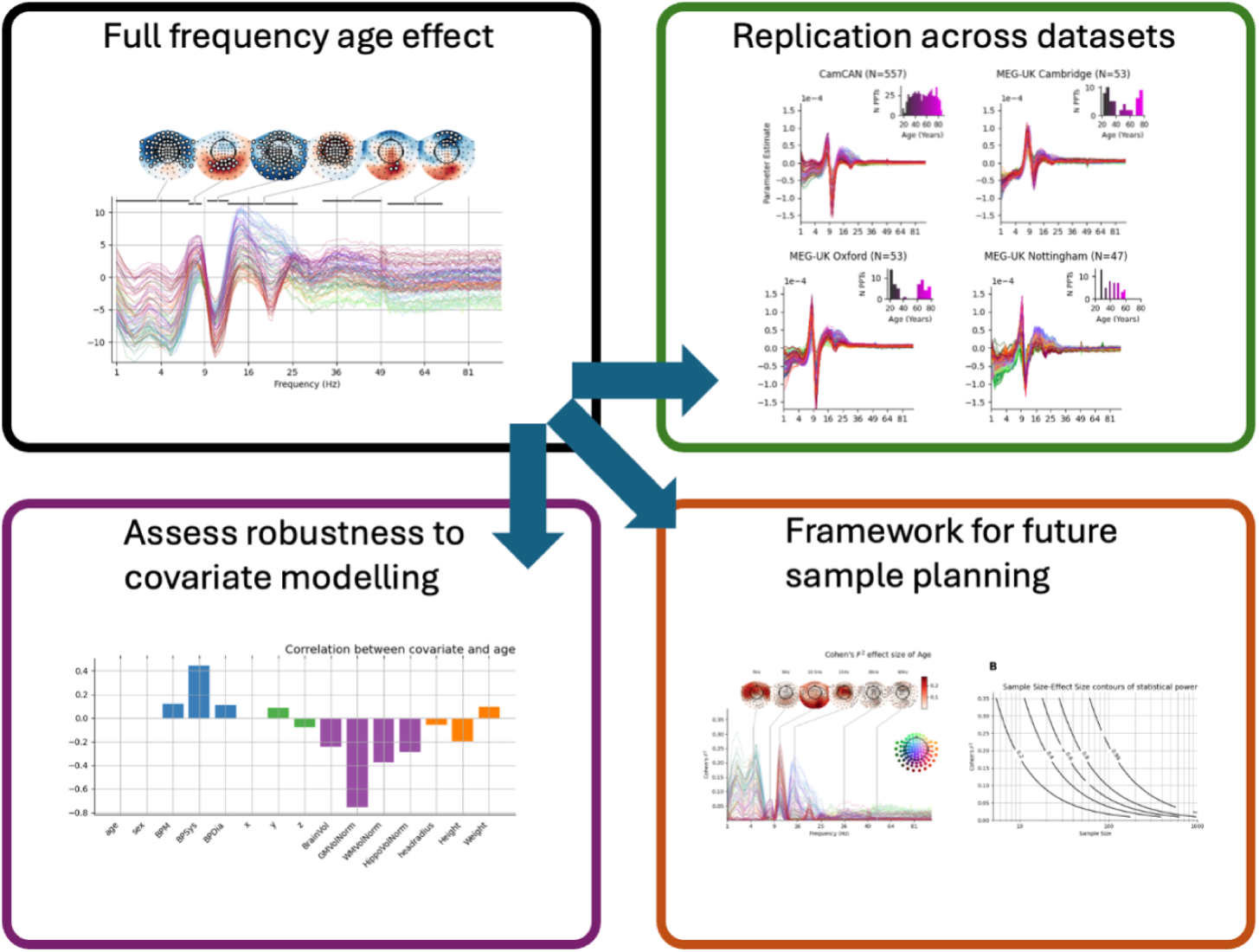

**Highlights:**

- There is a strong effect of ageing across the adult lifespan on neuronal oscillations across a range of frequencies and spatial locations.
- The effect is replicable across multiple open-access datasets.
- The effect is robust to a range of sources of inter-subject variability.
- The age effect has heterogeneous effect size across space and frequency with implications for future study planning.

## 1 Introduction

Ageing is a major risk factor for a wide range of major pathologies of the brain [Lopez-Otın et al., 2013] including neurodegenerative disorders [Planche et al., 2022] and stroke [Ellekjær et al., 1997, Feigin et al., 2014]. Changes in human brain electrophysiology are observable through changes in the power spectrum of electromagnetic fields measured by EEG and MEG. These approaches offer a direct perspective on temporal synchronization of neuronal activity [Babiloni et al., 2020]. This synchronization depends on the integrity of network connections and is known to degrade throughout the ageing process [Maestú and Ferńandez, 2020]. Such effects have been observed in the neuronal power spectrum of both MEG [Brady et al., 2020, Gomez et al., 2013, Hoshi and Shigihara, 2020, Rempe et al., 2023, Stier et al., 2023, Osipova et al., 2005, Hughes et al., 2019] and EEG recordings [Klimesch, 1999, Park et al., 2024, Turner et al., 2023, Zibrandtsen and Kjaer, 2021]. Non-invasive measures of brain electrophysiology are a highly promising approach to contribute toward the global aim of allowing more people to realise their potential across the full lifespan [World Health Organization, 2015].

Electrophysiological signatures of ageing across studies in the literature are diverse, or even conflicting. It can be challenging to reconcile differences across differing analysis choices, frequency bands, and brain regions [Donoghue et al., 2021]. For example, increasing age has a longstanding association with a reduction in posterior alpha band (7-13Hz) power [Klimesch, 1999]. Yet, the broader literature reveals that this effect varies in magnitude, and may even reverse direction, across brain regions, frequency bands, and analysis pipelines [Gomez et al., 2013, Hoshi and Shigihara, 2020, Lodder and van Putten, 2011, Park et al., 2024, Pathak et al., 2022, Quinn et al., 2024, Rempe et al., 2023, Scally et al., 2018, Zibrandtsen and Kjaer, 2021]. Credible scientific claims must be supported by evidence of replicability in new datasets and robustness to reasonable variations in analysis [Botvinik-Nezer and Wager, 2023, Nosek and Errington, 2020, Nosek et al., 2022], but our ability to integrate results across studies is hindered by the large variety of analytic choices used in the literature. This makes it difficult to synthesize findings from multiple datasets, either as a manual comparison or a formal meta-analysis.

Neuroscience researchers working on MRI facilitate comparability and meta-analyses by computing and distributing whole-volume statistical maps along with results of individual studies [Gorgolewski et al., 2015]. A similar approach is possible for spectral analysis of MEG/EEG, e.g. computing and sharing full frequency and whole-head estimates of critical effects without binning results into frequency bands or regions of interest. Features such as canonical frequency bands may be a useful guide for interpretation of results, but they can oversimplify the profile of the underlying spectrum and add a source of variability between studies that can prevent aggregation of results. Reporting of whole-head and full-frequency spectra of effect estimates would make it straightforward to aggregate across studies and eventually enable identification of sub-threshold effects that may be missed in single analyses but are consistent across studies. We argue that this approach provides a generalisable foundation that can support more complex analyses that have a frequency component (such as aperiodic slopes, burst detection, and dynamic functional networks) that require more researcher degrees of freedom.

Effect sizes are another essential tool for comparing and aggregating results across studies. They quantify the magnitude, or practical significance, of an estimated effect without reducing it to a hypothesis test with a binary outcome that might be dependent on practical factors such as sample size [Smith and Nichols, 2018]. Critically, effect sizes allow efficient use of data by supporting planning for future data samples through power analyses. Statistically underpowered studies are more likely to lead to incorrect conclusions and inaccurate estimates of effect sizes when all other aspects of experimental design and analysis have been conducted perfectly [Button et al., 2013, Cremers et al., 2017]. This is critical for neuroimaging analyses where different spatial, temporal or spectral components of the same signal feature can have different effect sizes [Baker et al., 2021]. Many guidelines on good practice in electrophysiology research encourage reporting effect sizes [Gross et al., 2013, Hillebrand et al., 2018, Baker et al., 2021] though this remains relatively uncommon in electrophysiology.

Here, we use a GLM-Spectrum [Quinn et al., 2024] approach to quantify and visualise age effects in the Cam-CAN dataset [Shafto et al., 2020]. The method uses multiple regression to model the relationships between multiple explanatory variables and the oscillatory power at each spatial location and frequency bin. The approach is expanded with between-subject effect sizes estimation in addition to replication on multiple datasets. An additional exploration examines how the age effect is changed when including other covariates in the model. The results are discussed in the context of the wider literature on age effects and lay the groundwork for rigorous evaluation and benchmarking of potential biomarkers for brain health in ageing.

## 2 Results

We first quantify the change in the MEG frequency spectrum across the adult lifespan using the Cam-CAN dataset in section 2.1. Section 2.2 explores the effect sizes of the adult lifespan results and Section 2.3 expands these approaches to explore quadratic ‘U’ shaped effects of age. Section 2.4 provides bootstrapped estimates of effect sizes to compute sample size guidance for planned replications of our results. We next demonstrate the reproducibility of these results across several MEG-UK datasets in Section 2.5 and their robustness across analyses that control for a range of potential confounding variables in Sections 2.6 and 2.7. Finally, we quantify the impact of including grey matter volume as a covariate on the effect size of the age effect in Section 2.8. A summary of all GLM analyses carried out in this work is available in supplemental section A.7.

### 2.1 A whole-head and full-frequency depiction of age-related change to the neuronal power spectrum

MEG data from Cam-CAN [Shafto et al., 2020] for 557 participants during eyes-closed resting-state were analysed. After data pre-processing and cleaning, a two-level GLM-Spectrum [Quinn et al., 2024] approach was used to characterise the average magnitude and the linear slope effect of age across the spectrum for each sensor. Figure 1A uses the estimated GLM-Spectrum coefficients to show the model prediction of how the spectrum changes across age. This shows a characteristic pattern that combines a variety of previously reported results into a single analysis. Importantly, these effects are continuous across the spectrum and would be challenging to objectively unpick without A full-spectrum approach.

**Figure 1.**
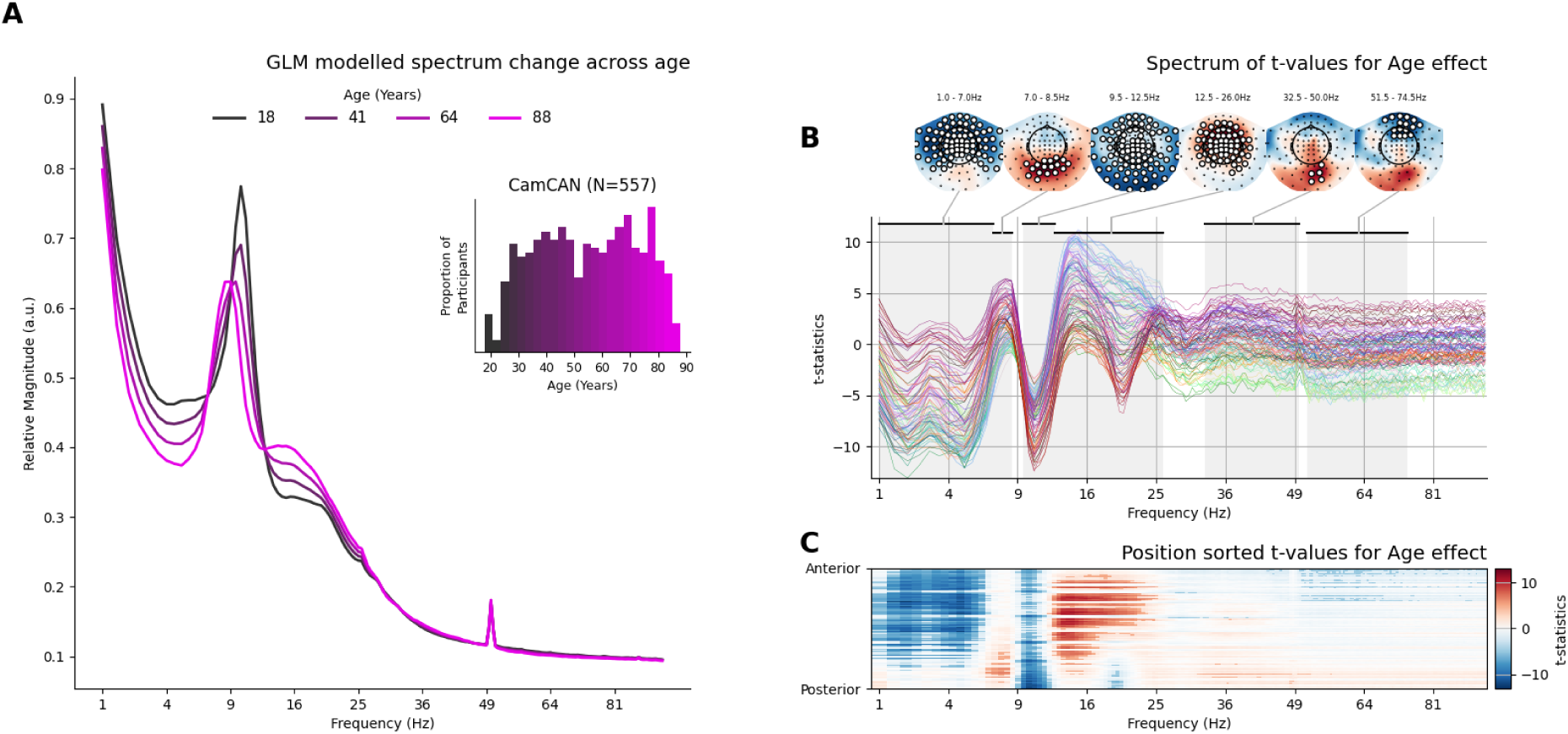
Effect of age on the relative magnitude spectrum across space and frequency. **A)** Model-predicted spectra (averaged across all sensors) for 4 equally spaced ages across the participant age range with inset histogram of participant ages within Cam-CAN. **B)** Spectrum of t-values quantifying the age effect across space and frequency. Non-parametric permutations with maximum statistics to control for multiple comparisons across sensors and frequency bins. While this permutation testing was not cluster-based, contiguous clusters of significant sensors were computed post-hoc for visualisation The largest 6 spatially and spectrally contiguous areas of statistically significant effects are highlighted in frequency by black bands at the top of the spectrum and highlighted in space by sensors marked with a white circle in the adjoining topography. **C)** A 2D frequency-by-space map of all statistically significant effects. Sensor-Frequency combinations that do not reach statistical significance have a faded colour scale. Blue regions indicate decreasing spectral power with age and red regions indicate increases.

The t-statistic spectrum (Figure 1B) is the test statistic for the linear effect of age at each point in frequency for each sensor. A positive t-value indicates that power in that frequency bin at that sensor position increases with age, and vice versa for negative t-values. The statistical significance of the t-statistic spectrum of the age-effect (i.e., rejecting the hypothesis that there is no age effect) was computed using non-parametric permutations with maximum statistics to control for multiple comparisons across sensors and frequency bins. While this permutation testing was not cluster-based, contiguous clusters of significant sensors were computed post-hoc for visualisation. Figure 1B shows the t-value spectrum of the age with the largest post-hoc clusters overlaid and Figure 1C shows the effect organised into a 2D image across space (anterior to posterior) and frequency as a statistical parametric map [Litvak et al., 2011]. The 2D image representation simplifies visualisation as it does not rely on colour-coding of sensor location. Critically, both the t-statistic spectrum and 2D image are simple visualisations of the age effect that allow statistically significant and sub-threshold effects to be shared and reused in the literature.

The maximum-statistic permutations testing results identified a broad set of effects, including six contiguous regions within the age-effect spectrum (Figure 1B). Firstly, older adults have lower spectral magnitude than young adults at low frequencies (*<* 7Hz). This is a spatially broad effect that peaks in central/frontal regions. This analysis does not separate different effects in low frequency bands such as delta and theta. Rather, they appear as a contiguous effect that spreads across the whole 1-7Hz range. Older adults have relatively high magnitude around 7-8.5 Hz, and younger adults have higher magnitude around 9.5-12.5 Hz. Both effects peak in posterior/occipital regions, with the negative age effect at higher frequencies spreading more widely across the sensor array. These effects are visible as distinct peaks with opposite direction (Figure 1B) in the GLM Spectrum of the age effect. Referring to the model predicted spectra (Figure 1A) suggests that this may reflect a slowing of the alpha peak frequency. Older adults have higher spectral magnitude between 12.5-26 Hz in central/motor regions corresponding to sensori-motor band activity. Finally, two effects are identified with centre frequencies above 30 Hz. Older adults have higher relative magnitude in posterior central sensors at frequencies between 32.5 and 50 Hz and have lower relative magnitude in fronto-central regions in the gamma region between 51.5 and 80 Hz (frequencies above 100Hz were not included in the model).

We have completed several control analyses to support these findings. Firstly, we have explored the qualitative correspondence between alpha peak frequency and the two effects we identified within the canonical alpha range (see supplemental section A.1). Secondly, the overall pattern of findings is consistent in an equivalent source space analysis using LCMV beamforming and parcellation (see supplemental section A.2). Finally, we have repeated the analyses with and without ICA denoising and find that the overall spectral profile is very similar. Effects in low-frequencies, low-alpha and high-gamma are increased with application of ICA, whilst high-alpha and beta remain unchanged, and low-gamma effects are reduced (see supplemental section A.3).

**Figure 2.**
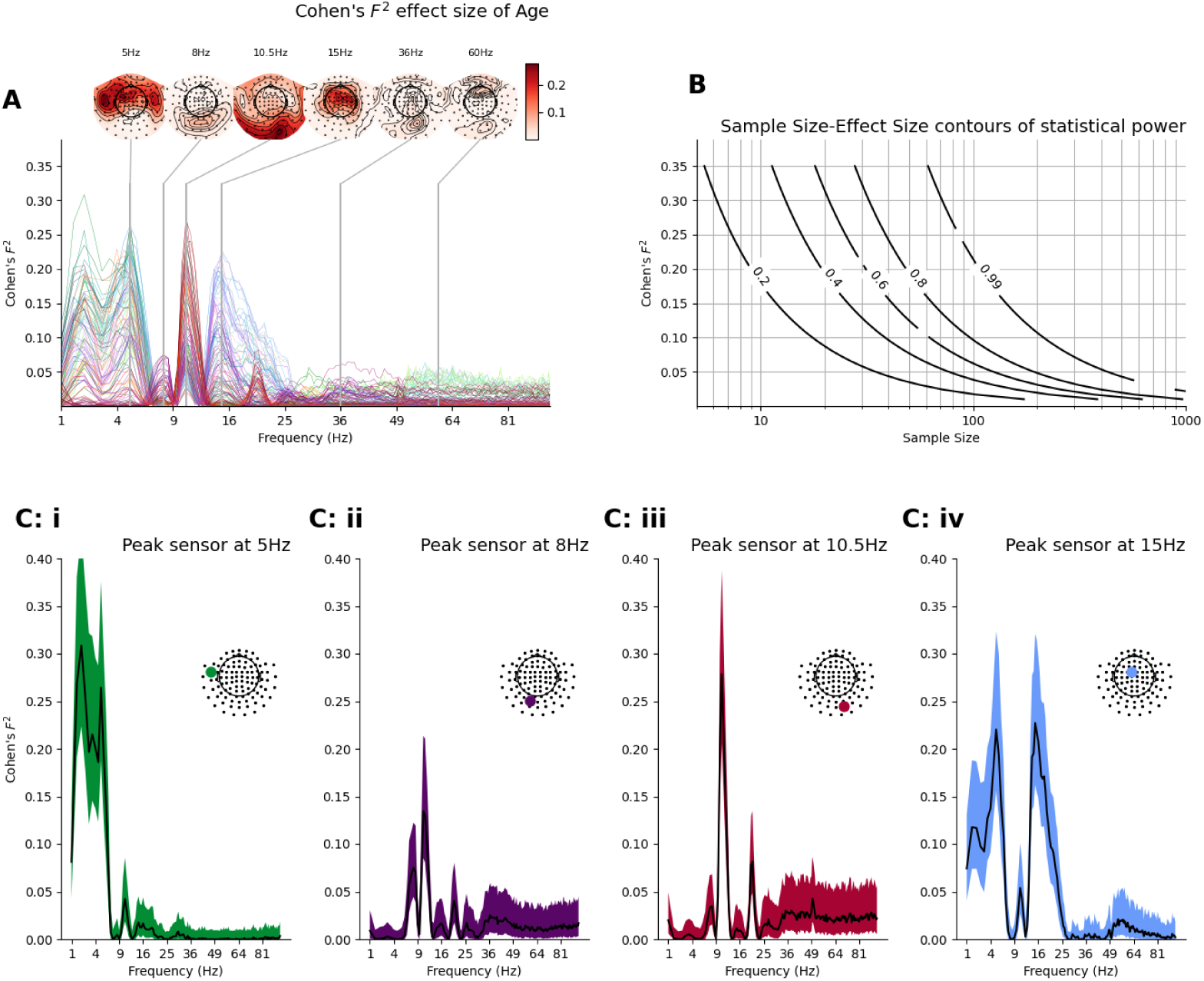
Cohen’s *F*^2^ effect size for the linear effect of age **A)** Cohen’s *F*^2^ effect size for age across space and frequency. Each line is a sensor, with a colour matched to the sensors in the topography shown in the inset. The six topographies show spatial distributions across the sensor array at the peak frequency of the six significant effects identified in Figure 1B. **B)** Contours showing the relationship between effect size and sample size for five different experimental power levels. As sample sizes get larger, there is sufficient power to reliably detect smaller effect sizes. **C)** Effect size spectra with bootstrapped 95% confidence intervals at the peak sensor for the four largest effects identified in Figure 1B.

### 2.2 Effect size of age is variable across space and frequency, with consequences for sample size planning

Neuroimaging analyses, such as the results in Figure 1, typically focus on testing hypotheses about the presence of effects in brain imaging data, not estimating their magnitude. These hypothesis tests are sample-size dependent, so that practically insignificant effects may become significant in sufficiently large datasets. Effect sizes estimate the strength of a phenomenon or relationship, independently of sample size. They better reflect an effect’s importance in a clinical or behavioural context [Reddan et al., 2017] and can be used as a basis for power analysis (i.e. sample size planning) to inform future research. In this analysis, the effect size of the age regressor within the GLM-Spectrum was computed using Cohen’s *F*^2^ [Cohen, 1988, Selya et al., 2012] for each sensor and frequency bin in the analysis. Cohen’s *F*^2^ corresponds to the proportion of variance in the data that is uniquely explained by an explanatory variable (i.e., regressor) of interest within the context of a multiple regression model.

The spectrum of effect sizes for the age regressor (Figure 3A) indicates the strength or practical impact of ageing on the spectrum. The effect size spectrum contains local peaks that correspond to the effects highlighted in section 2.1. The largest effects with local peaks at 5Hz, 8Hz, 10.5Hz, 15Hz, 36Hz, and 60 Hz have close spatial and spectral correspondence to extrema in the t-spectrum in Figure 1B. Low frequency, upper alpha and beta range results show effect sizes peaking between *F*^2^ = 0.2 to *F*^2^ = 0.3, indicating that ageing had a relatively large practical impact on the spectrum in these bands. In contrast, though they achieved statistical significance in the hypothesis tests (Figure 1), lower alpha and the gamma results had effect sizes around *F*^2^ *≤*0.075, indicating a relatively small practical impact of ageing on the spectrum. These effect sizes can be used to construct sample size estimates to inform future study planning. Power contours [Baker et al., 2021] across different effect sizes and sample sizes can provide a visual guide to the number of participants required to achieve a given statistical power in a future replication of the present analyses (Figure 3B).

**Figure 3.**
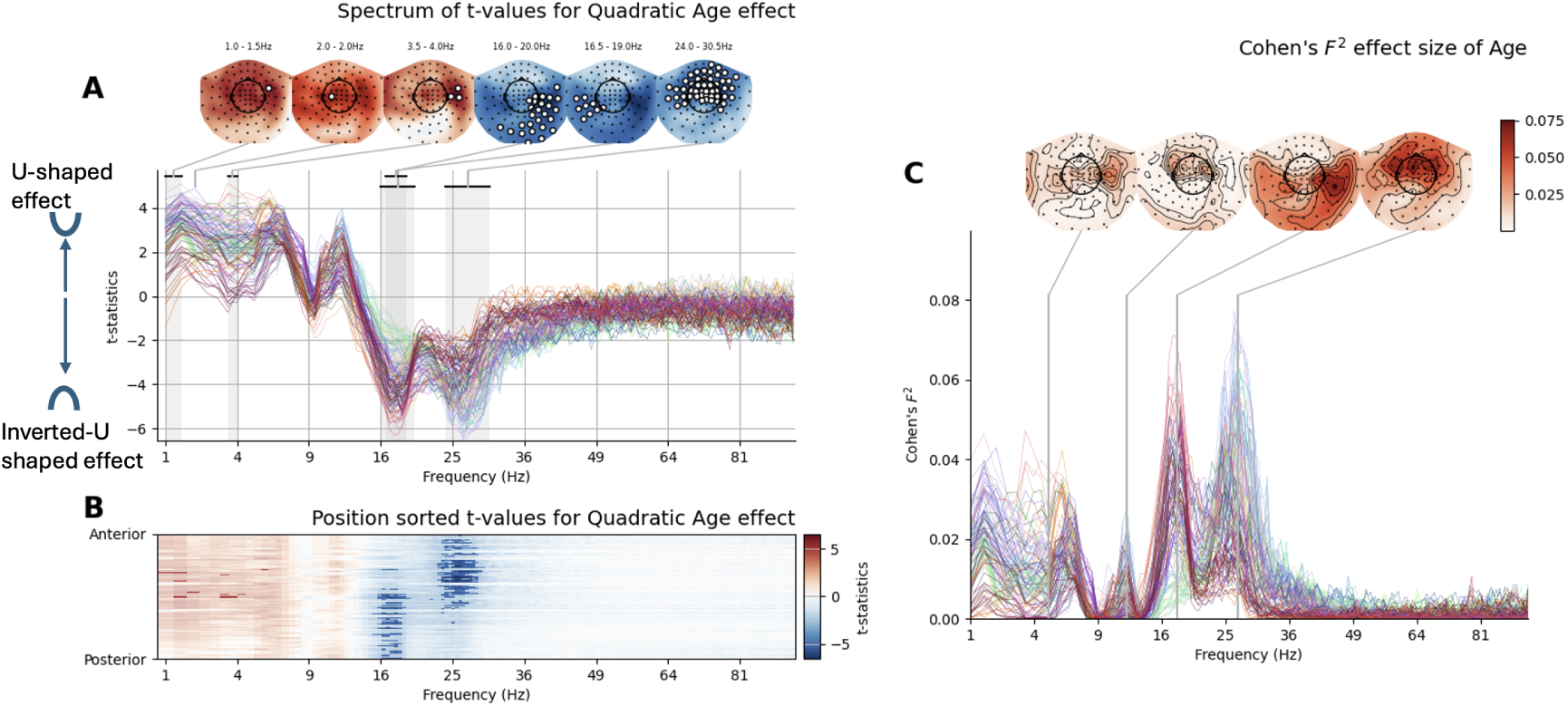
Quadratic effects of age on the neuronal frequency spectrum. **A)** The spectrum of t-values for the quadratic effect of age for all frequencies and channels. Positive values indicate a ‘U’ shaped effect and negative values indicate an ‘inverted-U’ shaped effect. Shaded regions and sensors indicated on the topomaps indicate statistical significance. **B)** A 2D frequency-by-space map of all statistically significant effects. Sensor-Frequency combinations that do not reach statistical significance have a faded colour scale. Blue regions indicate inverted-U shaped effects of age and red regions indicate U shaped effects. **C)** The spectrum of effect sizes for the quadratic effects of age.

It is important to acknowledge that effect sizes are estimates themselves and can vary in their precision depending on the dataset and analysis. To quantify this variability, bootstrapped confidence intervals were computed for each sensor and frequency pair. The computed confidence limits of the effect size estimates vary substantially across sensors and frequencies (Figure 3C shows the effect sizes with confidence intervals at the peak sensor for the first four significant effects).

### 2.3 Quadratic effects of age

The linear effect of age is a convenient and simple regression model. However it makes a strong assumption that change with age is uniform across the whole age range. A second group-level GLM was computed with an additional regressor to quantify quadratic effects of age, which have been reported in the ageing literature [Gomez et al., 2013, Rempe et al., 2023, Stier et al., 2023]. The spectrum of t-values for the quadratic age predictor (Figure 3) shows significant effects for a U-shaped change with increasing age in the low frequency (1-5Hz) range in central sensors. Significant inverted-U shaped effects are present in two frequency ranges in the beta band. A low-frequency beta effect is present in occipital and temporal sensors between 16Hz and 20Hz, whilst a second high-beta effect is present in central sensors between 24Hz and 30.5Hz. The low-frequency and low-beta effects overlap in space and frequency with linear effects, suggesting that the low-frequency change with age has both a linear increase with age and a U-shaped component, whilst the low-beta effect has a decrease with age and an inverted-U shaped component. The high-beta effect does not overlap in frequency with any of the reported linear effects. The effect sizes for quadratic effects range between Cohen’s *F*^2^ values of 0.033 for low-frequency to 0.076 for high-beta and are generally lower than the effect sizes for linear change.

### 2.4 Sample size planning for effects of age on the neuronal power spectrum

We use the 95% confidence intervals around the effect sizes to make recommendations for future sample sizes for future samples that plan to replicate these results. The observed power calculations have no bearing on the interpretation of the present results. Instead, they should be used as a general guideline for planning future studies. Table 1 gives a full summary of the peak statistics and future sample size range for the six age effects identified in Figure 1B. These results have implications for future sample planning for resting-state electrophysiology studies of ageing. Using the upper bound of the sample size estimates as a conservative estimate, the linear changes with age that have relatively large effect sizes would have well powered replications with sample sizes of around 50-60 participants. However, the smaller linear effects and all the quadratic effects would require samples of 200 or more participants to have the same probability of detecting the effect if it is indeed present (Table 1). This indicates that study samples should be planned with the smallest effect of interest in mind and that ageing effects in different frequency bands may not all be well powered within the same sample.

**Table 1.** Summary of significant spatio-spectral regions containing an age effect in Cam-CAN. The estimated sample sizes are the range of sample sizes needed to have an 80% chance of detecting the effect in a future sample, assuming that both the effects here are true and that the effect sizes are well estimated. Sample size ranges are computed from bootstrapped 95% confidence intervals for effect size estimate. The lower bound of the 95% confidence intervals can be taken as a smallest effect size of interest. The sample size forecasts are not informative about the present results and serve only as a guide for future study planning.

| Descriptive Label | Peak Freq (range) | Peak t-value | Peak Region | Peak Cohen's $F^2$ (95% CI) | Estimated Future sample for 80% Power |
| --- | --- | --- | --- | --- | --- |
| **Linear Effects** |  |  |  |  |  |
| Low Freq | 5Hz (1 - 7) | -12.11 | Frontal | 0.264 (0.187 - 0.375) | 26 - 51 |
| Low Alpha | 8Hz (7 - 8.5) | +6.44 | Occipital | 0.074 (0.044 - 0.122) | 79 - 218 |
| High Alpha | 10.5Hz (9.5 - 12.5) | -12.43 | Occipito-Parietal | 0.278 (0.205 - 0.387) | 25 - 47 |
| Beta | 15Hz (12.5 - 26) | +11.22 | Central | 0.227 (0.162 - 0.320) | 30 - 60 |
| Low Gamma | 36Hz (32.5 - 50) | +5.77 | Occipital | 0.06 (0.030 - 0.111) | 86 - 315 |
| High Gamma | 60Hz (51.5 - 74.5) | -5.37 | Frontal | 0.052 (0.024 - 0.098) | 97 - 396 |
| **Quadratic Effects** |  |  |  |  |  |
| Left Beta | 17 (16.5 - 17) | -4.62 | LH Occipito-Temporal | 0.033 (0.036 - 0.11) | 82 - 264 |
| Right Beta | 18 (16 - 20) | -6.26 | RH Occipito-Temporal | 0.071 (0.044 - 0.13) | 76 - 222 |
| Central High Beta | 27 (24 - 30.5) | -6.47 | Anterior Central | 0.076 (0.044 - 0.12) | 80 - 218 |

### 2.5 The spectral profile of age effects on brain electrophysiology is replicable across datasets

To explore the replicability of the ageing effect reported from the Cam-CAN dataset, the analysis was repeated on eyes-closed resting-state recordings from three smaller datasets from the MEG-UK database (https://meguk.ac.uk/database/). Details on the similarities and differences between the datasets are summarised in Table 2. Replicability was quantified by the correlation of the whole spectral profile of the age effect between datasets. We have only completed the replication analysis with the linear effects of age as the quadratic effects from Cam-CAN have smaller effect sizes that indicate that the MEG-UKI datasets may not be able to detect these effects based on their sample size.

**Table 2.** Summary of significant spatio-spectral regions containing an age effect in Cam-CAN. The estimated sample sizes are the range of sample sizes needed to have an 80% chance of detecting the effect in a future sample, assuming that both the effects here are true and that the effect sizes are well estimated. Sample size ranges are computed from bootstrapped 95% confidence intervals for effect size estimate. The sample size forecasts are not informative about the present results and serve only as a guide for future study planning.

| Analysis & Descriptive label | Age Effect Size | Age Effect Size with GGMV | Change in age effect size from including GGMV |
| --- | --- | --- | --- |
| **Effect size estimates** |  |  |  |
| Low freq (5 Hz) | 0.26 (0.19 - 0.38) | 0.11 (0.065 - 0.18) | -0.15 (-57%) |
| Low alpha - 8 Hz | 0.075 (0.043 - 0.13) | 0.013 (0.0016 - 0.039) | -0.048 (-83%) |
| High alpha (10.5 Hz) | 0.28 (0.20 - 0.40) | 0.12 (0.080 - 0.20) | -0.16 (-57%) |
| Beta (15 Hz) | 0.23 (0.16 - 0.32) | 0.078 (0.045 - 0.13) | -0.152 (-66%) |
| Low gamma (36 Hz) | 0.06 (0.031 - 0.11) | 0.071 (0.034 - 0.14 ) | +0.011 (+15%) |
| High gamma (60 Hz) | 0.052 (0.024 - 0.096) | 0.044 (0.015 - 0.10) | -0.008 (-15%) |
|  | **Sample with 80% Power for Age** | **Sample with 80% Power for Age including GGMV** | **Change in sample for including GGMV** |
| **Sample size planning** |  |  |  |
| Low freq (5 Hz) | 25 - 52 | 54 - 148 | +30 to +98 |
| Low alpha (8 Hz) | 76 - 225 | n.s. | n.s. |
| High alpha (10.5 Hz) | 24 - 47 | 49 - 120 | +25 to + 73 |
| Beta (15 Hz) | 30 - 60 | 73 - 216 | +43 to +156 |
| Low gamma (36 Hz) | 85 - 304 | 67 - 285 | -18 to -19 |
| High gamma (60 Hz) | 100 - 401 | n.s. | n.s. |

Overall, the spectral profile of the ageing effect is highly replicable across MEG datasets acquired on different systems and across different facilities. We identified high correlations between the spectral profiles of the age parameter estimates, inferential statistics, and effect sizes in each of the four datasets (Figure 4). In particular, the spectral profiles for the parameter estimates and t-statistics were strikingly similar across datasets (Pearson’s R between 0.67 and 0.97 for parameter estimates and between 0.76 and 0.92 for t-statistics), suggesting a high level of replicability. The spectral profile of effect sizes was also positively correlated across datasets but to a lesser extent (Pearson’s R between 0.53 and 0.84). Maximum and minimum t-statistics for Cam-CAN are substantially larger in magnitude than in the smaller MEG-UK datasets. This is an expected consequence of the larger sample size of Cam-CAN on null-hypothesis test statistics such as t-values.

**Figure 4.**
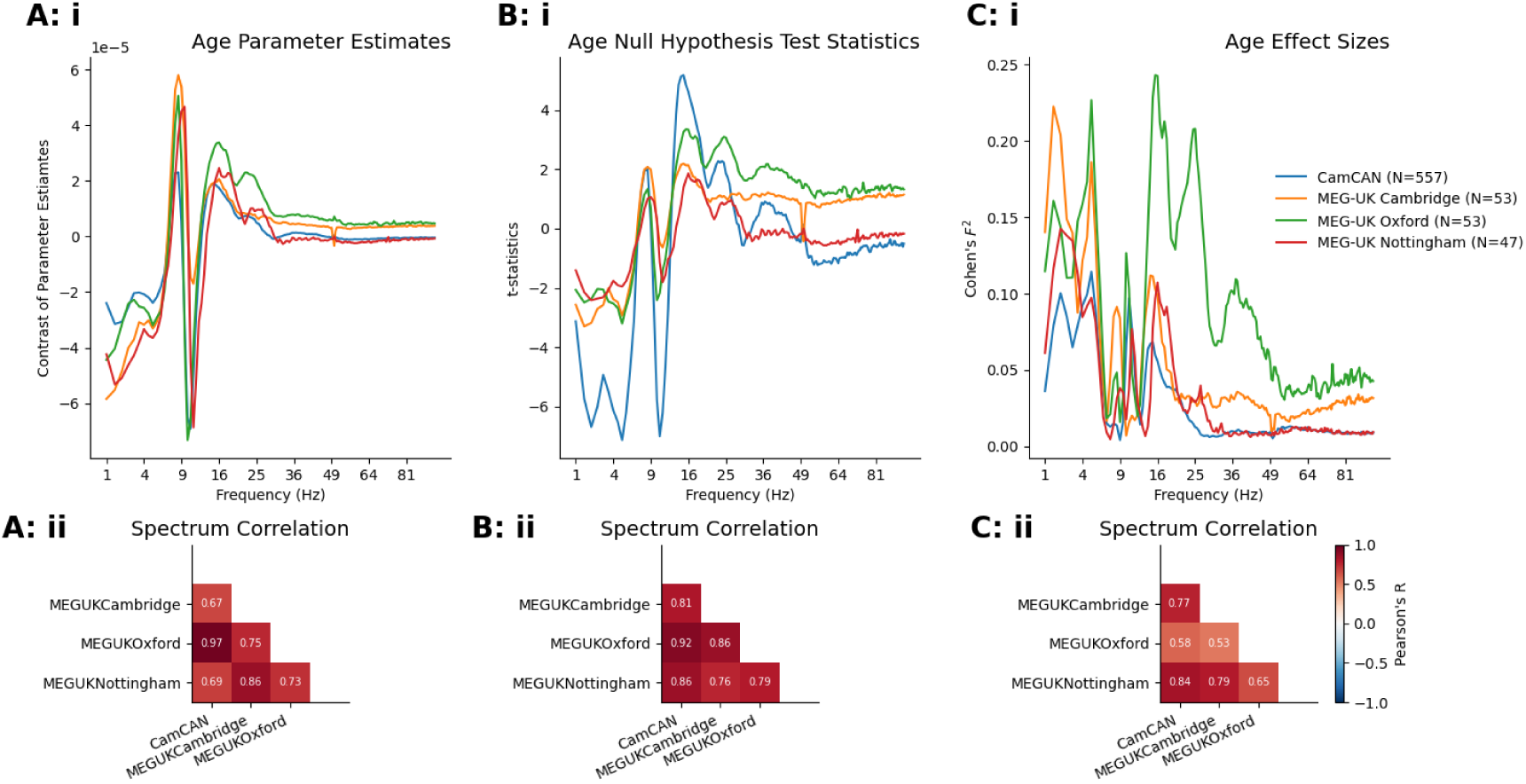
Replicability of the parameter estimates, t-statistics, and effect sizes of ageing on the neuronal power spectrum. **Ai)** Spectrum of parameter estimates across frequency (averaged over all sensors) for each of the four datasets. **Aii)** Correlation matrix indicating similarity of the sensor-averaged frequency profile of the age effect between datasets. **Bi & Bii)** As A for null hypothesis test statistics. **Ci & Cii)** As A for effect sizes.

The topography of age-related change is qualitatively reproduced across all datasets (Figure 5A) with some variability in the extent and spread of effects in specific datasets. Whilst the lower frequency effects are highly consistent, there is more variability in the higher frequency effects (*>* 30 Hz), which can change direction across participants (Figure 5B, C D). The smaller data samples have more variable estimates for Cohen’s *F*^2^ when compared to Cam-CAN. Whilst the interpretation of effect sizes doesn’t rely on reaching a predetermined significance threshold, they do depend on having enough data to be considered reliable. This is reflected in the greater variability in effect sizes seen in the smaller data samples. The overall profiles of Cohen’s *F*^2^ are similar across datasets, yet the peak effect size in the low-frequency and beta ranges can still vary by a factor of two or three between data samples. This may arise from relatively poor estimates of the population-level variability from smaller data samples.

**Figure 5.**
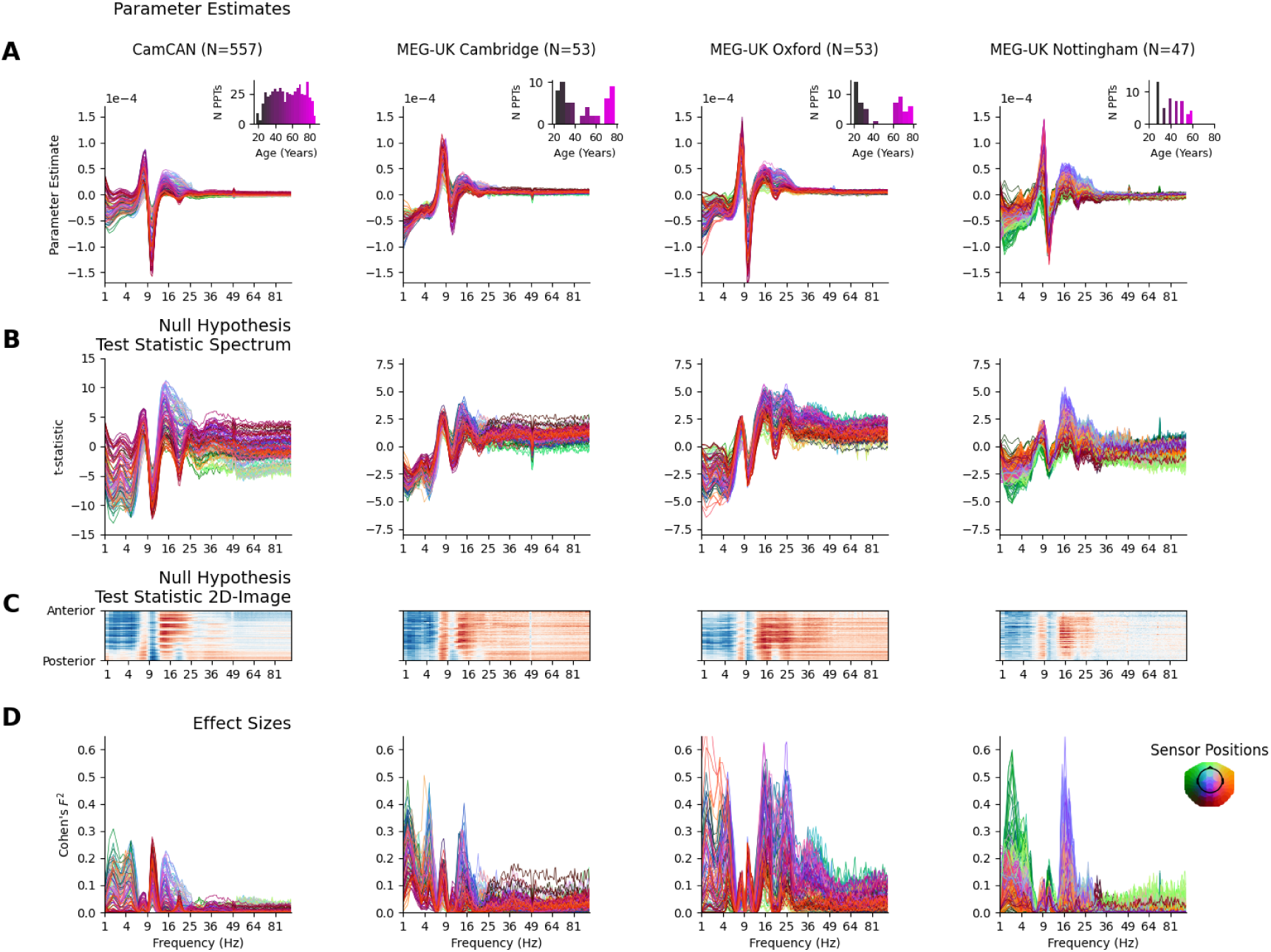
The spatial and spectral profile of the effect of age on resting-state brain electrophysiology is replicable across datasets. **A)** GLM-Spectrum parameter estimates quantifying the age effect across space and frequency for the four datasets in the replicability analysis with distribution of participant ages shown in the inset figure. The four datasets share the key features highlighted in Figure 1. **B)** As A) for t-statistics testing the hypothesis that the parameter estimate of the age effect is different to zero. **C)** As B) but visualised as a 2D image in which each row indicates a single sensor with their y-axis position sorted by spatial location on the anterior-posterior axis. **D)** as A) for Cohen’s *F*^2^ effect sizes for the age regressor in the GLM-Spectrum.

It is possible that more systematic differences in the participant sampling, recruitment, and data acquisition process could contribute to between-site differences. At present, our analyses can show that the core effects of ageing are replicable across several datasets, though we have not formally combined these datasets into a single analysis. Formal methods for data harmonisation, such as ComBat [Johnson et al., 2006], could correct for additive and multiplicative differences in data scaling across sites to improve site comparisons.

### 2.6 The robustness of the age effect to head position correction

The results so far have looked at age effects on the relative spectral magnitude of each frequency compared to the sum across frequencies. Whilst the absolute spectral magnitude is simpler to interpret in terms of underlying neuronal activity it can be confounded by the position of the participant relative to the MEG sensors (some sensors will be closer to neuronal source than others), and by inter-individual variability in this position (Gross et al., 2013). It is common to apply a correction or normalisation to sensor-space analyses to control for these effects, though this is an impactful decision which can change the profile of effects seen in the results.

The age effect on the absolute spectrum (Figure 6) presents a different pattern of results compared to the relative spectrum (Figure 1 & 3). Briefly, the beta effect remains preserved or even enhanced compared to the relative magnitude effect, whilst the alpha effects are attenuated, and the low-frequency effect is decreased or removed altogether (Figure 6A, B). There is a reduced spatial specificity in the age effects of absolute power. The largest post-hoc cluster of contiguous statistically significance for absolute power covers a broad range of frontal and temporal sensors and stretches the whole 1-95-Hz range of the spectrum (Figure 6C) whilst the next four largest clusters have relatively homogenous spatial patterns (Figure 6C). Spatial maps are positively correlated for all pairs of frequencies (Figure 6B), indicating that absolute scaling from head position might be making a large contribution to the overall effect.

**Figure 6.**
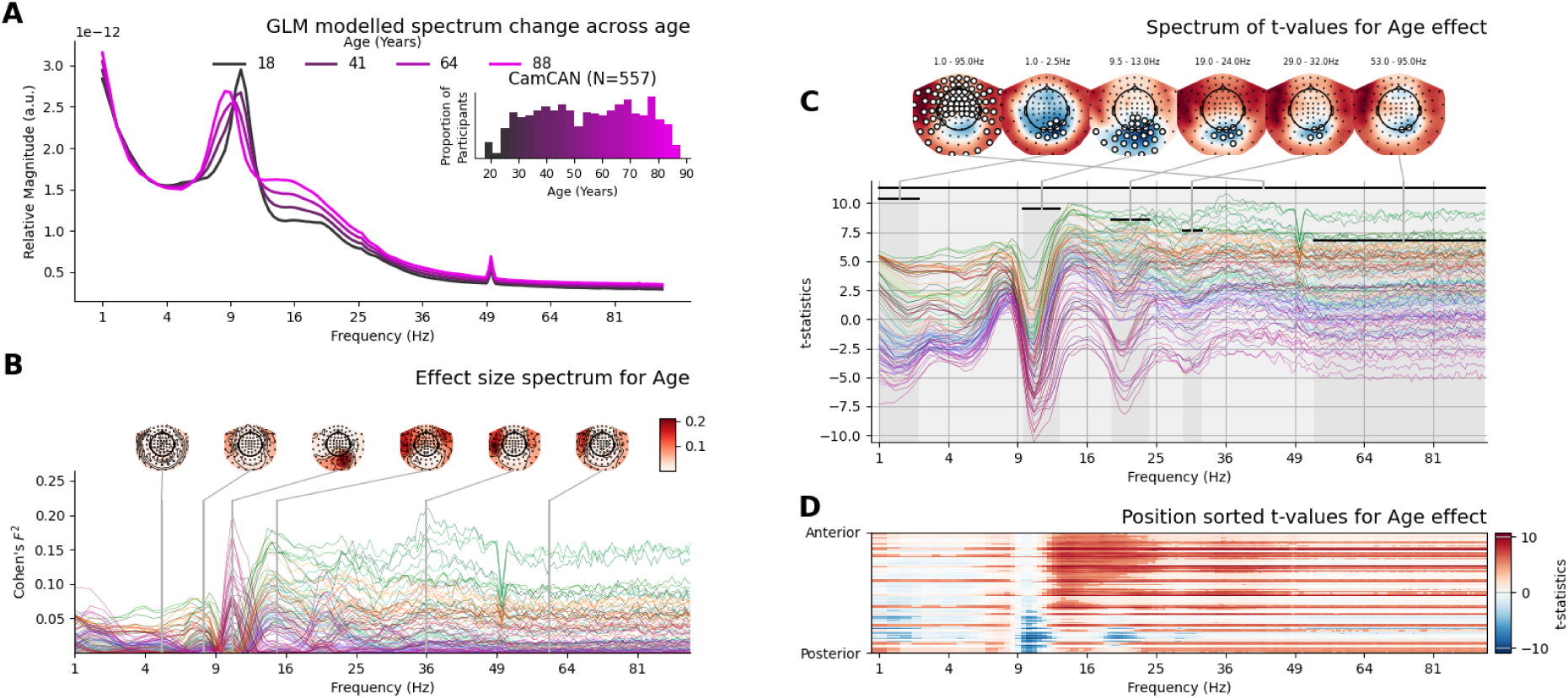
The age effect computed on the absolute magnitude of the power spectrum. **A)** Model-predicted spectra (averaged across all sensors) for 4 equally spaced ages across the participant age range with inset histogram of participant ages within Cam-CAN. **B)** The effect size for the age regressor computed on the absolute magnitude of the power spectrum. **C top** Spectrum of t-values quantifying the age effect across space and frequency. Non-parametric permutations with maximum statistics to control for multiple comparisons across sensors and frequency bins. While this permutation testing was not cluster-based, contiguous clusters of significant sensors were computed post-hoc for visualisation The largest 6 spatially and spectrally contiguous areas of statistically significant effects are highlighted in frequency by black bands at the top of the spectrum and highlighted in space by sensors marked with a white circle in the adjoining topography. **C bottom** A 2D frequency by space map of all statistically significant effects. Sensor-Frequency combinations that do not reach statistical significance have a faded colour scale. Blue regions indicate decreasing spectral power with age and red regions indicate increases.

A related decision involves rescaling data recordings to correct for differences in head position within (movement compensation) and between (transformation to reference position) data recordings. This is commonly applied using Signal Space Separation (SSS) with the maxfilter software [Taulu and Kajola, 2005, Taulu and Simola, 2006]. We additionally compared the age effect across absolute and relative spectral magnitude with three variants of SSS head-position correction (None, movement compensation, and movement compensation plus transformation to reference). The spectral profile of the absolute effect (without relative scaling) is relatively consistent across the three variants in SSS head position correction with the application of between recording transformation to reference head position creating the only noticeable difference (Figure Supplemental 17A, C & E). In contrast, the relative spectrum has the familiar spectral and spatial profile from Figure 1 and is unaffected by SSS head position correction (Figure Supplemental 17B, D & F). There is substantially greater spectral specificity in spatial profile of the age effect in relative power, indicated by greater variability in spatial correlations across frequency (Figure Supplemental 17G & H).

### 2.7 Between-participant covariates mediate the age effect differently across the spectrum

Next, we explored whether the age effect is robust when controlling for covariates other than age, i.e. putatively confounding explanatory variables. These alternative covariates may be correlated with age and may impact the estimated relationship between ageing and the observed neuronal power spectrum in complex ways. We selected a non-exhaustive range of alternative covariates split into five categories: demographic (sex), cardiac (heart rate, systolic and diastolic blood pressure), MEG data acquisition (head position in dewar), brain anatomy (brain volume, global grey matter volume, global white matter volume, hippocampal volume), and physiological (head radius, height, weight) factors. We do not distinguish between covariates representing neuronally relevant factors and those representing sources of non-neuronal variance.

The GLM-Spectrum was used to quantify the effect of each alternative covariate on the spectrum by fitting a separate GLM for each of the alternative covariates that only contained a regressor for the alternative covariate, alongside an intercept term. Brain volume, global grey matter volume (GGMV), systolic blood pressure, and sex had the strongest association with the observed MEG power spectrum, though the effect size varies strongly across frequency and space (Figure 5A & Supplemental Material).

Next, we compare the age effect in two models: the first includes an age regressor plus intercept, and the second model includes an age regressor, a single alternative covariate regressor, and intercept term. The difference in Cohen’s *F*^2^ effect size for age estimated in a simple linear regression model and the partial effect of age from a multiple regression model quantifies how robust the age effect is to the inclusion of the covariate. Including GGMV and systolic blood pressure strongly reduces the estimated age effect sizes, whilst inclusion of brain volume, white matter volume, or hippocampal volume leads to either increases and decreases in the observed age effect depending on the specific sensor and frequency bin (Figure 5B & supplemental material).

Covariates with low correlation to age generally did not modify the estimated age effect (e.g. head position, heart rate, and weight). In contrast, including covariates with high correlation can lead to starkly different influences in different brain regions and frequency ranges. For example, including GGMV GG or systolic blood pressure strongly reduced the estimated effect size of age across the brain. In contrast, inclusion of white matter volume or brain volume in the model equally increased and decreased the estimated age effect size, depending on the position and frequency being observed. Systolic blood pressure, white matter volume, and brain volume have comparable correlations with age, but the impact of that correlation on the estimates of the age effect is substantially different.

### 2.8 Isolating the component of the age effect on brain electrophysiology that is distinct from global reduction in grey matter volume

Global grey matter volume (GGMV) had the strongest mediating effect on the impact of ageing on the MEG spectrum from all the alternative covariates (Figure 7). Whilst there is overlap between age and GGMV in their effect on the MEG spectrum, each has a unique contribution that is distinct from the other. Modelling GGMV alongside age reduced the association between age and neuronal power spectra heterogeneously across different frequency bands and did not reduce the age effect to zero (Figure 8A and B).

**Figure 7.**
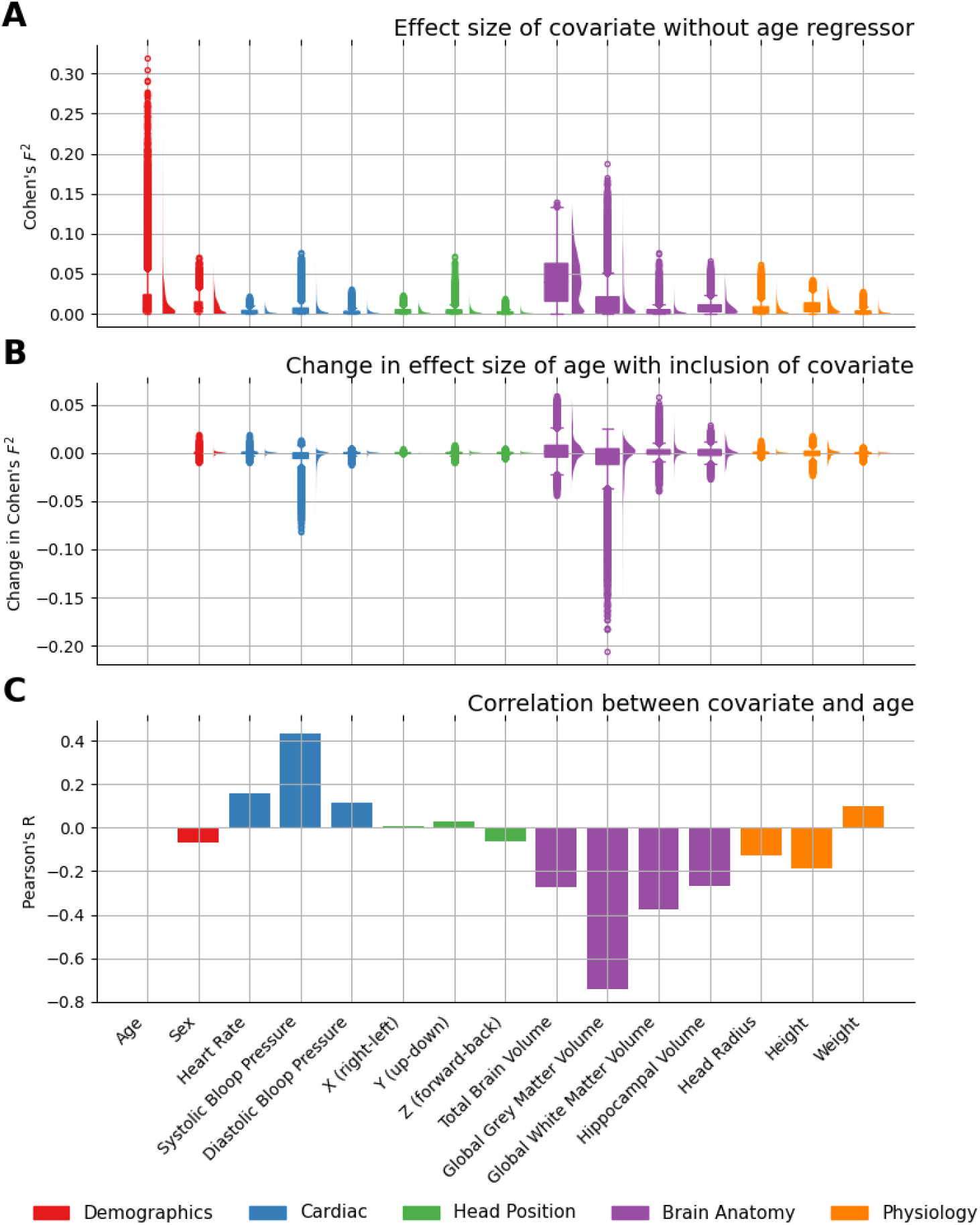
Covariate effect sizes and their impact on the ageing effect. **A)** Box and whisker plot with paired kernel density plot for the distributions of Cohen’s *F*^2^ effect size for each alternative covariate, when the alternative covariate is included as the only regressor in a GLM, along with an intercept term. The Cohen’s *F*^2^ distributions are collected over every single sensor-frequency pair from the sensor-space dataset (i.e. over 102 sensors and 189 frequency bins). Full GLM spectrum visualisations of the results are included in the appendix. For comparison, the same is shown for the age using a GLM that contains regressors for age and an intercept. **B)** Change in the Cohen’s *F*^2^ effect size of age between 1) a GLM that includes an age regressor plus intercept, and 2) a model that includes an age regressor, a single alternative covariate regressor and intercept. This is shown for each alternative covariate in turn. Age is excluded from this panel. **C)** Pearson’s correlation coefficient quantifying the univariate linear relationship between age and each covariate. Age and sex are excluded from this panel.

**Figure 8.**
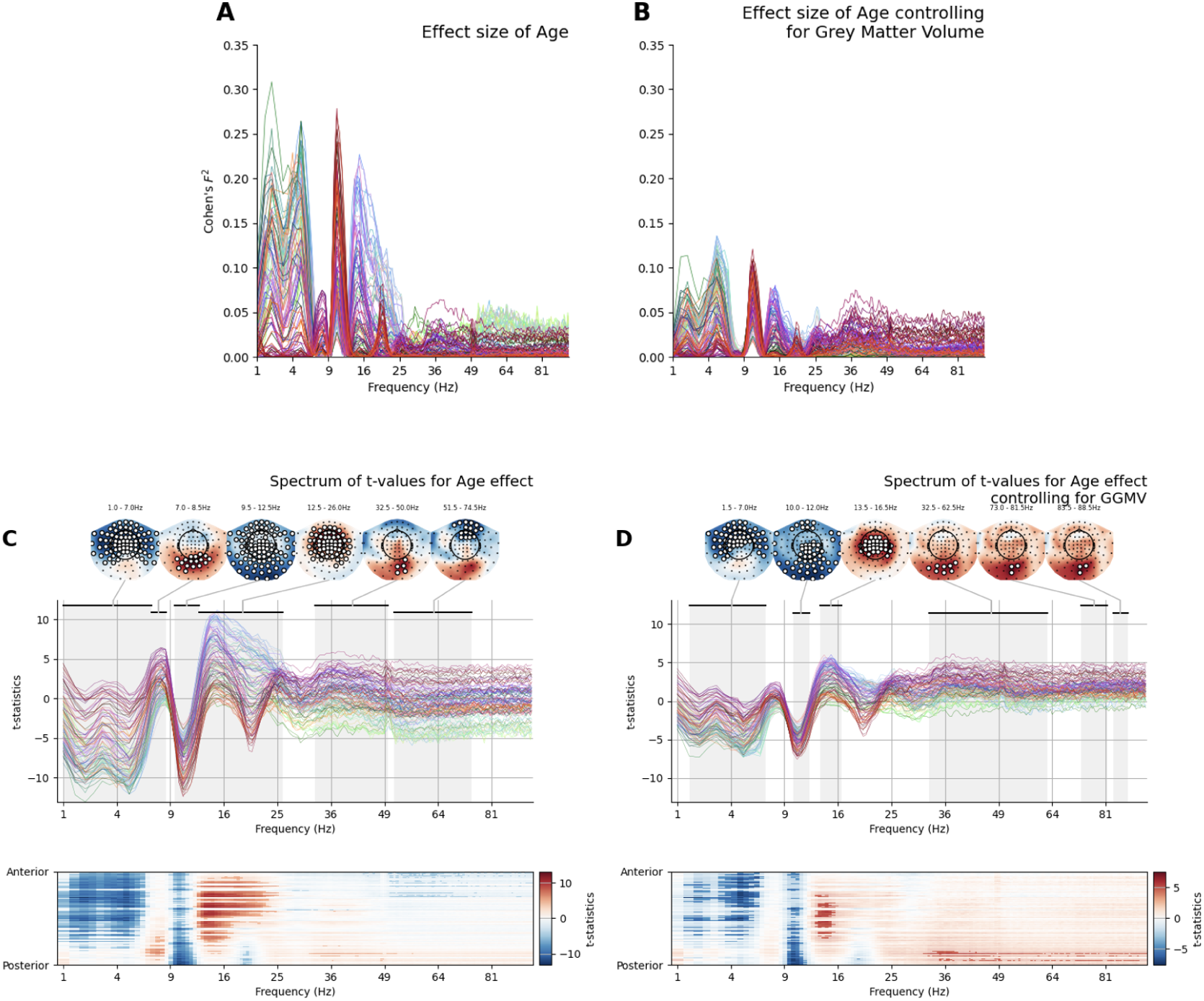
The age effect is reduced heterogeneously across space and frequency by including GGMV in the model. **A)** The Cohen’s *F*^2^ spectrum for age using a GLM that contains regressors for age and an intercept. Replicated from Figure 3. **B)**as A) when a Grey Matter Volume regressor is added to the GLM. Accounting for grey matter volume broadly reduces the effect size of age across the spectra, but some moderate effects remain. **C)** Spectrum of t-values quantifying the age effect across space and frequency replicated from Figure 1. Non-parametric permutations with maximum statistics to control for multiple comparisons across sensors and frequency bins. While this permutation testing was not cluster-based, contiguous clusters of significant sensors were computed post-hoc for visualisation The largest 6 spatially and spectrally contiguous areas of statistically significant effects are highlighted in frequency by black bands at the top of the spectrum and highlighted in space by sensors marked with a white circle in the adjoining topography **D)** As C) for the age effect in a model that includes a global grey matter volume covariate. Most effects are reduced in size with smaller statistically significant regions surviving the post-hoc clustering procedure.

It is important to note that the correlation between age and GGMV does impact the interpretation of the GLM results, but does not prevent the model fit. We explore the model validation and diagnostics in detail in Supplemental section A.6. In brief, the model is able to separate the unique contribution of age and GGMV. However, the correlation between factors leads to an inflation in the standard error of the estimates. A hypothesis test on these estimates is valid though the inflated variance increases type-2 error by reducing our ability to detect partial effects in the data when they are truly present.

Whilst most of the age effect-size spectrum was reduced by inclusion of GGMV, the extent of this reduction varied clearly across the six spatio-spectral regions that showed significant simple linear age effects (See Table 2). Five of the six regions showed a reduction in peak effect size (between −15% and −83%), with a single effect showing a small increase (Low gamma range +15%). The regions with the largest initial age effect sizes (before GGMV inclusion) showed the greatest reductions when controlling for GGMV. Despite these reductions, the remaining effect sizes in these regions were still larger than the original effect sizes observed in the regions with initially smaller effects.

These results have a substantial practical impact on study planning. A sample that appears to be well powered for estimating an effect when considering the effect in isolation may in fact be underpowered when covariates and confounds are accounted for in the regression model. Using the present results from Cam-CAN, a study looking to replicate our findings of about the component of the age effect that is linearly distinct from GGMV will require a larger sample than one replicating the age effects alone (Table 2). Most critically, two (low alpha and high gamma) of the six identified age effects drop below the significance threshold when accounting for GGMV (Figure 8). This suggests that a large part of the electrophysiological effects in these bands cannot be linearly separated from global structural change with ageing.

## 3 Discussion

A whole-head and full-frequency profile of the ageing effect is characterised and shown to be replicable across datasets. Six distinct ageing effects are identified, and the combination of linear parameters across the spectrum can represent more complex features such as alpha peak frequency. The corresponding effect sizes demonstrate that age impacts the spectrum heterogeneously and that the feature of interest must be considered when sample size planning for future experiments. This effect is robust to head position correction, but the spatial profile changes substantially between relative and absolute spectral power. Finally, the age effect is differentially modified by different factors, such as grey matter volume, brain volume, head size, and physiological factors. Individual variability in grey matter volume mediates the association between age and neuronal power spectra, leading to a reduction in the estimated effect size of age across the whole spectrum. Critically, the impact of including a grey matter volume covariate differentially impacts different brain regions and frequency bands, indicating that some regions have a strong and unique signature of ageing that reflects functional/synaptic changes over and above grey matter atrophy.

Overall, by analysing four independent datasets including 600 participants, we demonstrated that there is a replicable and robust effect of ageing on MEG power spectrum across the whole-head and a range of frequencies. These effects are well-powered at moderate data samples, replicable across different datasets and scanner types, and are reasonably robust to modelling additional anatomical, physiological, or acquisition factors as covariates.

### 3.1 Full frequency spectra of age effects synthesise diverse results across the literature

The profile of results across the spectrum of age effects replicates and simplifies a range of results across previous literature. This is done without pre-specification of frequency bands or sensors of interest and in a manner that is conducive to future study planning and meta-analyses. Here, we briefly review the core age effects from this analysis in relation to a representative sample of published results on the effects of healthy ageing on the resting-state EEG and MEG spectrum.

**Low-frequency power decreases with age**. This result is broadly consistent with literature demonstrating that ‘theta’ power decreases with healthy ageing [Beese et al., 2017, Cummins and Finnigan, 2007, Rempe et al., 2023, Vlahou et al., 2014] and may be associated with decreases in cognitive performance [Finnigan and Robertson, 2011]. One critical difference with this literature is that the full-frequency age profile identifies a single ‘low frequency’ age-effect between 1-7 Hz and does not distinguish between the canonical delta (2-4 Hz) and theta (4-7 Hz) oscillations. A further difference is that we only see the low-frequency decrease in power in the relative spectrum (Figure 1). Computing the age effect from the absolute spectrum resulted in no changes across age at low frequencies (Figure 6).

A combination of age-related decreases in lower frequencies with increases in higher frequencies (notably the beta range) contributes to a flattening of the aperiodic component of the power spectrum [Aggarwal and Ray, 2023, Cesnaite et al., 2023, Dave et al., 2018, Voytek et al., 2015]. This flattening is proposed to reflect an age-related increase in neural noise affecting age-related working memory performance [Voytek et al., 2015]. Future research accounting for flattening of the 1/f-type component in the full frequency spectrum approach can better separate whether the low frequency changes relate to oscillations or the aperiodic component of the spectrum.

**Reconciling conflicting reports of the ageing effect on alpha power**. The literature exploring how ageing changes alpha power is heterogeneous. Papers that report results in resting alpha power, either from a canonical band or from an individual peak frequency, include reports of a variety of contrasting age effects. This includes positive correlations with age [Rempe et al., 2023, Stier et al., 2023], negative correlations [Thuwal et al., 2021, Lodder and van Putten, 2011, Medrano et al., 2025, Park et al., 2024], both positive and negative effects separated by space [Hoshi and Shigihara, 2020, Pathak et al., 2022], or null results when correcting for individual frequencies and aperiodic slopes [Scally et al., 2018, Merkin et al., 2023] (see supplemental section A.1 more detailed summary). There is broad variability in methodological approaches, which could account for the variety of results. Stier et al. [2023] suggest that analyses in sensor or source space may lead to different effects. Critically for our work, this variability also prevents formal aggregation of results and meta-analyses that could clarify the picture.

We have proposed that by taking a step back, and tolerating a reduction in sensitivity, we can map out the whole-head whole-frequency structure of the age effect with minimal researcher degrees of freedom and bias. Investigating age effects as a complete spectrum shows that two contrasting age effects on alpha magnitude coexist in close proximity in space and frequency: an increase with a small effect size in central occipital sensors around 7-8.5 Hz and a decrease with a large effect size across a broad set of occipital, temporal and frontal sensors between 9.5-12.5 Hz. Different data samples and different data analysis choices, particularly the selection of regions of interest, source reconstruction, sensor normalisation or correction of aperiodic components or might emphasise one effect or the other in each analysis. Whilst we have not conclusively explored all possible variants of these analyses, we have provided a framework that would allow future studies to perform formal comparisons and meta-analyses to resolve this bottleneck.

**Relationship between effects on alpha power and alpha individual frequency**. The two effects identified in the present analysis may combine to represent a decrease in power and frequency of a single alpha peak (seen qualitatively in Figure 1A). This change in alpha peak frequency is highly replicable [Cesnaite et al., 2023, Dustman et al., 1993, Sahoo et al., 2020, Scally et al., 2018, Pathak et al., 2022, Zibrandtsen and Kjaer, 2021] and is a highly predictive spectral marker of ageing [Stier et al., 2024]. Decreases in alpha peak frequency have been linked to a decline in cognitive performance in healthy ageing [Cesnaite et al., 2023, Finley et al., 2024] and MCI [Garćes et al., 2013, Ĺopez-Sanz et al., 2016, Puttaert et al., 2021].

This compelling evidence for a shift in a single peak is complicated by strong evidence for presence of multiple alpha peaks within individuals [Lodder and van Putten, 2011, Chiang et al., 2011, 2008, Klimesch, 1999] with distinct generators and functional relevance [Sokoliuk et al., 2019]. A complex pattern of changes in power, frequency and spatial distribution likely underlies age-related change in alpha oscillations. Future work will need to explore all three features at the individual level to clearly illuminate the change.

**Linear increase and inverted-U effects in the beta band** The literature reports an increase in low beta power with age [Gomez et al., 2013, Heinrichs-Graham and Wilson, 2016, Heinrichs-Graham et al., 2018, Hübner et al., 2018, Koyama et al., 1997, Rempe et al., 2023, Stier et al., 2023, Veldhuizen et al., 1993, Xifra-Porxas et al., 2019]. We have observed inverted-U shaped quadratic effects of age in two frequency ranges in the beta-band, a posterior low-beta component (centred around 18Hz) and an anterior high beta component (centred around 25Hz). This is broadly consistent with reports of quadratic ageing effects in the beta band in the literature [Gomez et al., 2013, Rempe et al., 2023, Stier et al., 2023], though to our knowledge, our results are the first to suggest a separation of effects into different parts of the beta range. These spectrum changes may be associated with age-related changes in underlying bursting dynamics [Brady et al., 2020, Power et al., 2023].

In addition to changing in the resting-state, ageing is associated with increases in movement-related beta desynchronisation [Bardouille and Bailey, 2019, Heinrichs-Graham and Wilson, 2016, Heinrichs-Graham et al., 2018, Xifra-Porxas et al., 2019, Rossiter et al., 2014]. Changes in post-movement beta rebound have been reported as well [Bardouille and Bailey, 2019, Xifra-Porxas et al., 2019]. The increased resting-state beta activity may require increased downregulation in order for older adults to perform movements [Heinrichs-Graham and Wilson, 2016].

**Novel gamma band changes with age**. There are relatively few publications exploring changes in effects of ageing on resting-state gamma band activity. Two EEG studies report increases in gamma power in older adults compared to younger adults [Aoki et al., 2022, Jabes et al., 2021]. The frequency band of these effects is consistent with our finding though the EEG results indicate a frontoparietal and temporal shift whereas our gamma increase occurs in occipital sensors. There is larger literature on task related changes in gamma with age which indicates that gamma responses to visual gratings decrease in power and frequency with age [Murty et al., 2020, Kumar and Ray, 2023].

**Mediation of age effects by grey matter volume**. There are substantial reductions in grey matter volume and cortical thickness across the adult lifespan [Frangou et al., 2022]. We show that association between age and the resting spectrum is mediated by reductions in grey matter volume but that this effect is heterogeneous across space and frequency. Our results show that accounting for GGMV reduces the effect sizes of the age effects below 30Hz by 57% to 83% (Figure 8; Table 2). This reduction replicates results showing that structural brain changes are associated with widespread changes in spectral power [Stier et al., 2023]. Our results are limited by the use of global grey matter volume as a blunt measure of grey matter change, grey matter decline in ageing occurs heterogeneously across the brain [Frangou et al., 2022] and oscillatory changes are likely linked to this pattern [Mahjoory et al., 2020].

### 3.2 Observed power analysis for future study planning

The observed power and sample size calculations in this paper should not be used to help interpret the present findings, or as a basis to speculate about what sample size would be required for a non-significant result to pass the significance threshold. Rather they are intended solely to guide decisions about the sample sizes of future studies in an accessible manner and to highlight the potential for this approach in analyses of neuronal power spectra.

We concur with Lenth [2001] who highlight that “Sample size planning is often important, and is nearly always difficult”. Most recommendations about sample size planning indicate that *a priori* or theoretically informed effect sizes should be used as observed effect sizes are influenced by sampling error [Hoenig and Heisey, 2001]. Single estimates of observed effect sizes can be extremely noisy and will not necessarily reflect the ‘true’ underlying population effect. Moreover, using the single observed estimate simply recapitulates the information already present in the p-value of the test [Hoenig and Heisey, 2001].

As a result, we emphasise the bootstrapped 95% confidence intervals of the effect sizes to give a sense of the uncertainty around our estimates. The upper and lower bound of these confidence intervals are used in the observed power analysis as an example to guide future study planning. Until such a time when the field reaches consensus about what the smallest effect size of interest would be for electrophysiological power spectra, we propose that researchers can use the lower bound of the 95% confidence interval as the smallest effect size of interest for a replication. Working with multiple covariates and a requirement to correct for multiple comparisons both decrease the statistical power of a neuroimaging dataset [Cremers et al., 2017] so it is critical to be transparent about which effects are observable in our data. Confidence interval based sample projections from observed effect sizes must be handled with care but are a useful tool in a field where sample size plans are frequently based on resource constraints or heuristics. Projections can be supported with further sensitivity power analyses that explore a realistic range of expected effect sizes given the context of the planned project. Many excellent guides have been written to inform principled sample size planning in behavioural [Lakens, 2022] and neuroimaging contexts [Mumford, 2012].

We propose the following steps as a practical guide for researchers looking to plan a data sample with a reasonable chance of correctly identifying a particular effect.

1. **Define research question and identify previous results:** Your research question must be defined in advance and well specified, there should be relevant data or literature that can be used to guide your decision.
2. 2. **Define the smallest effect size of interest, and the decision criterion for the sample decision:** It is critical to define the parameters of how you will make your sample size decision ahead of time. We recommend considering what the ‘smallest effect size of interest’ [Anvari and Lakens, 2021] would be for your question. This is specific to your question and is about more than statistical significance. What effect size would indicate that there is a *practically meaningful* effect to for future literature to consider?
3. **Estimate effect sizes to inform your decision:** Either by aggregating reported statistics from the literature, or by dedicated processing of previous data, compute an estimate of the effect size. Effect sizes are only estimates so it is important to compute confidence intervals around your estimate to get a measure of variability.
4. **Compute power/precision curves assuming these results:** Using the estimated effect sizes, compute power curves [Baker et al., 2021] that visualise the relationship between effect size, statistical power and sample size.
5. **Select the sample size that meets your pre-defined criteria:** Using your definitions from step 2, and when considering the whole power curve, make a decision about what sample size would give you a reasonable chance of replicating the effect of interest
6. 6. **Make note of any differences or deviations from this plan during your data collection:** There are many practical reasons why your planned sample might not match the data acquired in practice. Such deviations from a plan are ok, but should be acknowledged and any mitigating steps explained [Lakens, 2024].
7. 7. **Document your process for inclusion in a preregistration or publication:** Include details on how a sample size decision was made in your research outputs including preregistrations, preprints, and publications. These details will be useful for future researchers to understand your process and to implement their own.

### 3.3 Limitations

Our approach promotes an exploratory and unbiased approach to quantifying the age effect on neuronal power spectra which is intended to complement more focused analyses. This has the benefit of reducing researcher’s degrees of freedom and of being broadly generalisable. These come at the cost of a loss in specificity and in sensitivity. Our approach does not specifically quantify features derived from the power spectrum such as alpha-peak frequency or the aperiodic component of the spectrum. These features are mixed into our full spectrum estimates but not directly quantified. Thus they can be challenging to interpret from our approach. Secondly, the mass-univariate approach suffers from a potential loss in sensitivity compared to results that aggregate across spatial or spectral regions that contain consistent results. Where a region or frequency band of interest can be supported from the literature, an approach focusing on a single region has the benefit of reduced noise by averaging estimates from a larger range of observations. Finally, models that consider the whole shape of the spectrum [Donoghue et al., 2021] would also be able to combine information across a range of frequencies rather than depending on a single frequency bin for each estimate. These models have the additional benefit that their parameters are often directly interpretable as features of interest, such as spectral slope or peak frequency.

Whilst we have explored the impact of several analysis decisions on the profile of the ageing effect, we have not completed an exhaustive search of the processing options available to electrophysiology researchers. We have demonstrated that the age effect is robust to SSS based head position correction and to a broad range of covariates but modified by choice of relative or absolute power and inclusion of grey matter volume as a covariate. A broad range of analysis options are still to be explored, notably including the impact of different source reconstruction methods beyond the single example presented here. This is not an exhaustive exploration of the garden of forking paths [Gelman and Loken, 2013] and future analysis can broaden this scope through approaches such as multiverse analysis methods [Clayson et al., 2021, Dafflon et al., 2022].

Both absolute and relative power measures are used throughout the literature, but there is little consensus about their interpretation [Sandre and Troller-Renfree, 2026]. We focus on relative power for most of our results. By normalising each participant’s power spectrum by the sum across frequencies, we found that the results gained specificity in frequency band and reduced concern about wide inter-subject differences in overall variance. Though this improved some analyses, relative power has important drawbacks. In particular, it can reduce sensitivity to effects that are broadly distributed across the spectrum and uses arbitrary scaling rather than meaningful physical units. We support calls in the literature to report both relative and absolute power measures [Rempe et al., 2023, Sandre and Troller-Renfree, 2026] as they contain complementary insights.

We have compiled results from a broad range of whole-head and full-frequency models to provide a comprehensive overview of the linear age effect. However, due to the complexity of the data, further analysis is still needed to explore various options. We have only explored the linear effects of age in these analyses, whilst there is evidence for quadratic age effects in power spectral features [Brady et al., 2020, Rempe et al., 2023]. Our current analysis pipeline may not adequately capture variability in individual data recordings by neglecting to incorporate first-level variance components into the group model. A mixed modelling approach would provide a more comprehensive and robust framework for analysing these data. Finally, we present results based on spectral magnitude rather than power or log-power as a compromise between suitability for Gaussian linear modelling and intuitive presentation of findings [Quinn et al., 2024].

### 3.4 Building towards robust and reproducible effects of age in brain electrophysiology

We propose that computing and sharing unthresholded whole-head and full-frequency statistical spectra of key effects can facilitate comparability between studies. In addition, and the spectra of ageing effect sizes can support formal meta-analyses and planning of future sample sizes. We propose that this simple approach can provide a robust and replicable foundation stone on which clinical applications and more advanced analysis methodologies can be built and validated.

Similar approaches have been successful in facilitating the synthesis and crosstalk of results across a diverse literature in MRI [Gorgolewski et al., 2015]. To date, meta-analyses of spectral effects of ageing have been limited to features within specified frequency bands, such as alpha band peak frequency [Freschl et al., 2022] or power [Lejko et al., 2020]. These analyses are currently limited by variable and often incomplete reporting of results in addition to cumbersome decisions about how to combine results across heterogeneous channel sets, spatial regions, and frequency bands. Best practice reporting guidelines [Gross et al., 2013] and formal reporting standards, such as COBIDAS-MEEG [Pernet et al., 2018] improve transparency, but there is still a need to share outputs in a form that can readily support synthesis across studies and establish consensus about core effects

### 3.5 Future extensions

A clear future extension for this work is to formally incorporate estimates of within-subject variability into a mixed-effect model. These powerful models would enable modelling of both fixed effects and random effects, allowing researchers to account for variation within individuals over time and between individuals. Linear Mixed Models also provide improved approaches for handling missing observations and unbalanced designs, making them especially useful for longitudinal and hierarchical data. A second expansion of this work could use Generalised Additive Mixed Models (GAMMs) to model non-linear relationships using smooth functions while also accounting for random effects. This may allow for more realistic representations of complex patterns of change across age. Linear Mixed Modelling comes with a substantial increase in researcher degrees of freedom and can be challenging to implement and report accurately [Meteyard and Davies, 2020]. We have used fixed effects modelling in this work in line with our objectives of maintaining generalisability and minimising researcher degrees of freedom.

### Conclusion

Overall, we have demonstrated that ageing effects on MEG power spectra are robust and replicable across multiple datasets. Different brain regions and frequency bands respond differently to ageing, with some having a strong unique signature of ageing that is dissociable from global anatomical changes. This is a necessary step towards adoption of electrophysiological measures in a healthcare setting.

## 4 Methods

### 4.1 Datasets

Four open-access cross-sectional datasets with participants covering a broad age range were analysed as part of this manuscript.

**Cam-CAN**. Data used in the preparation of this work were obtained from the Cam-CAN repository (available at http://www.mrc-cbu.cam.ac.uk/datasets/camcan/), [Taylor et al., 2017, Cam-CAN et al., 2014]. **MEG-UK Oxford, Cambridge & Nottingham** Eyes closed resting-state MEG recordings were acquired in four MEG laboratories in the United Kingdom as part of the wider MEG-UK consortium. The Oxford and Cambridge data was acquired on a 306 channel VectorView system whilst the Cardiff and Nottingham data was acquired on a CTF-275 channel system.

### 4.2 MEG Data Preprocessing

All MEG data pre-processing was carried out using MNE-Python [Gramfort, 2013] and OSL-ephys [Quinn et al., 2022, van Es et al., 2025] using the OSL batch pre-processing tools. For the Cam-CAN data, we proceeded with analysis on post-maxfilter processed data provided by the Cam-CAN team. Briefly, the data were processed using AA [Cusack et al., 2015] with automatic bad channel detection (limited to 7 channels) with the origin set to the centre of a sphere fitted to the individual’s Polhemus headshape points. Maxfilter signal-space separation was performed with the temporal extension enabled (temporal window of 10 second and correlation threshold of *r* = 0.98). Head position was continuously estimated and compensated for during periods where the HPI coils were on. After maxfilter processing head positions were translated into a default head position defined as a point relative to each individual’s origin in a head coordinate frame. Data with the full maxfilter processing and with the head position translation or movement compensation were extracted from the Cam-CAN database. An equivalent pipeline was implemented in OSL-Ephys and applied to the data from the MEG-UK datasets from Oxford and Cambridge. Files from the MEG-UK Nottingham dataset were processed with third-order gradiometry applied.

The data from each recording was first converted from its raw format into MNE-Python Raw data objects. The first 35 seconds of continuous data were cropped out, and the remaining data were bandpass filtered between 0.25 and 150 Hz using an order-5 Butterworth filter. 50-Hz line noise was suppressed using the ZapLine method [de Cheveigńe, 2020] as implemented in the MEEGKit toolbox. Two passes of ZapLine were applied to account for minor peaks around the 50 Hz artefact. One pass was centred on 49 Hz and the second on 50 Hz. Bad segments were identified by segmenting data into 2-second chunks and using the generalised-extreme studentized deviate [Rosner, 1983] algorithm to identify outlier (bad) samples with high variance across channels. Bad segment detection was applied to both the raw time series and the differential of the time series and separately to the Magnetometers and Gradiometers. An average of 2.82% of data samples were marked as ‘bad’ (standard deviation: 2.54%, min: 0.0%, max: 15.52%) equivalent to an average of 15.04 seconds per recording. Bad channels were then identified using the G-ESD routine to identify outliers in the distribution of variance per channel over time. The data were then resampled to 250 Hz to reduce space on-disk and ease subsequent computations.

Independent Component Analysis (ICA) denoising was carried out using a 64-component FastICA decomposition [Hyvarinen, 1999] on the MEG channels. This decomposition explained an average of 99% of variance in the sensor data across datasets. Artefactual components relating to eye movements or the heart rate were automatically identified by correlation with the simultaneous EOG and ECG channels. ECG artefacts were identified using cross-trial phase statistics [Dammers et al., 2008] and an automatic threshold based on the sample rate of the data as implemented in the mne.preprocessing.ICA.find bads ecg function in MNE Python. Between 0 and 3 EOG components were rejected in each dataset, with an average of 0.99 (standard deviation: 0.79) across all datasets. EOG artefacts were identified by correlation with the HEOG and VEOG channels with a threshold set to *r* = 0.35 as implemented in the mne.preprocessing.ICA.find bads eog function in MNE Python. Between 0 and 5 ECG components were rejected in each dataset, with an average of 2.25 (standard deviation: 0.84) across all datasets. The continuous sensor data were then reconstructed without the influence of the components labelled as artefacts.

To retain consistent dimensionality across the group, any bad channels were interpolated using a spherical spline interpolation [Perrin et al., 1989] as implemented in MNE-Python. The head position of the participant in device coordinates was extracted from each dataset.

This pipeline was repeated for both the SSS processed data with and without movement compensation and head position standardisation.

The same pipeline was applied to the VectorView data from the Oxford and Cambridge sites of the MEGUK dataset. A very similar pipeline was applied to the CTF data from the Nottingham site of dataset MEGUK dataset, except for omitting the SSS processing in favour of 3rd order gradiometry and in selection of planar gradiometer channels.

### 4.3 MRI Data Processing

All MR data were processed using the FMRIB Software Library [Woolrich et al., 2009].

T1-weighted MR scans from 804 individuals were analysed. Images were reoriented to MNI space, cropped, and bias-field corrected. FMRIB’s Linear Registration Tool, FLIRT [Jenkinson and Smith, 2001, Jenkinson et al., 2002] was used to register to standard space before brain extraction was performed using BET [Smith, 2002]. Tissue-type segmentation (grey matter, white matter and CSF) was conducted using FMRIB’s Automated Segmentation Tool, FAST [Zhang et al., 2001]. Subcortical structure volumes (hippocampus, amygdala, etc) were derived using FMRIB’s Integrated Registration and Segmentation Tool [Patenaude et al., 2011].

Total head volume was computed by extracting a volumetric skull mask using the FSL BETSurf tool and filling in the gap left by brain extraction. The total number of voxels within the head and brain was computed with fslmaths. The overall brain volume was normalised by the total head volume.

The voxel count for each tissue-type and subcortical structure was extracted and normalised using the individual’s total brain volume (computed by FAST) to compute a percentage. The final MRI metrics were: Total head volume, Brain volume (as a proportion of head volume), Global Grey Matter Volume (grey matter volume as a proportion of total brain volume), and Hippocampus volume (summed across left and right hemispheres as a proportion of total brain volume).

### 4.4 Source Reconstruction & Parcellation

First, a structural MRI (sMRI) was used to coregister the MEG data. This involved extracting surfaces for the outer skin, outer skull and inner skull from the sMRI using FSL, then aligning the Polhemus headshape points to the outer skin surface, as well as aligning the sMRI and Polhemus fiducials. Note, 31 out of 643 subjects from Cam-CAN were excluded at this stage due to missing sMRIs.

Following the coregistration, a single-shell forward model was computed. The preprocessed sensor-level data were source reconstructed using a Linearly Constrained Minimum Variance (LCMV) beamformer [Hillebrand et al., 2005, Van Veen and Buckley, 1988]. This involved projecting the sensor-level data onto an 8-mm dipole grid inside the inner skull. The forward model and sensor-level data covariance matrix were used to compute the LCMV beamformer. The covariance matrix was estimated using the entire recording for a subject and regularised to 60 components using principal component analysis (PCA) rank reduction.

Using the dipole time courses from source reconstruction, parcel time courses were calculated using an anatomical parcellation with 52 regions of interest (binary) [Kohl et al., 2023]. The first principal component across dipoles assigned to a parcel was used to generate the parcel time courses. Following this, the parcel time courses were orthogonalised using a symmetric multivariate correction for spatial leakage [Colclough et al., 2015]. Finally, a stochastic search algorithm was used to align the sign of each parcel time course across subjects by matching to a (median) template subject.

### 4.5 GLM-Spectrum - First level

A GLM-Spectrum [Quinn et al., 2024] was used to provide a statistical estimate of the MEG spectrum in the presence of covariates and confounds. A Short-Time Fourier Transform (STFT) was computed from the preprocessed sensor time series from each dataset using a 2-second segment length, a 1-second overlap between segments, and a Hanning taper. The 2-second segment length at the sample rate of 250 Hz gives a resolution of around 2 frequency bins per unit Hertz in the resulting spectrum. The short-time magnitude spectrum is computed from the complex-valued STFT and the frequency bins ranging between 0.1 and 100 Hz taken forward as the dependent variable in the first-level GLM-Spectrum for that dataset.

The GLM design matrix for each dataset is specified with five regressors. A single constant regressor models the intercept of the data, whilst two zero-mean parametric regressors model the effect of changing heart rate and a linear trend in time throughout the recording. The fourth and fifth regressors were non-zero mean parametric regressors modelling the effect of variance in the V-EOG channel and bad segments. These final regressors acted as confound variables that removed variance that can be linearly explained by the artefact sources from the mean term.

A set of simple contrasts isolated the main effect of each of the five regressors. The first-level GLM-Spectrum model parameters were estimated using the Moore-Penrose Pseudo-Inverse [Penrose, 1956] to compute beta-spectra. Finally, the beta-spectra were weighted by the contrasts to compute cope-spectra which were carried forward to the group-level analyses.

### 4.6 GLM-Spectrum - group-level effect of age

A group-level design matrix was constructed with two regressors, one constant intercept term, and one parametric regressor containing z-standardised participant age. The input data were the first-level average magnitude spectrum from each participant. The model coefficients were computed using the Moore-Penrose Pseudo-Inverse before contrasts and t-statistics were computed. Non-parametric permutation statistics were used to compute the statistical significance by row-shuffling the age regressor. Multiple comparisons were controlled by taking a maximum-statistic across all sensors and frequency bins. Contiguous clusters of significant effects were identified to simplify the final visualisation of the results. Cohen’s *F*^2^ was computed for the Age regressor for all sensors and frequency bins.

A second group-level design matrix was constructed to explore quadratic effects of age. This design contained three regressors, the intercept, a z-standardised age regressor and a z-standardised age regressor raised to a power of 2 to quantify quadratic components. The model fit, validation and statistical assessment of the quadratic predictor of this model was performed in the same way as the linear effect in the previous model.

### 4.7 GLM-Spectrum - group-level effect between subject covariates

Eleven potential between-subject covariates were collated for group analysis and are summarised in Table 2. To explore the individual effect of each covariate, a separate two-regressor model containing a constant term and a covariate was fitted for each covariate in turn. Non-parametric permutation statistics were used to estimate statistical significance for each of the covariates in these simple models. The effect size of each covariate in the simple models were computed using Cohen’s *F*^2^ statistics.

**Single Regressor Models:** The core model contained two regressors, one constant term and one z-transformed age regressor. This model was computed twice on different preprocessing versions of the Cam-CAN dataset, once using the full first-level GLM-Spectra and once without the additional covariates (Linear trend, EOG, ECG, and bad segments) in the first level.

**Age plus quadratic age models:** To move beyond linear change and explore U and inverted-U shaped effects of age, we fitted a group model with three-regressors. One constant term, one z-transformed age regressor, and one z-transformed quadratic age regressor (*age − mean*(*age*))^2^.

**Age plus covariate models:** the effect of each covariate on the overall effect of age was explored by fitting a second set of models that contain a constant term, the age covariate, and each individual other covariate in turn. Statistical significances and effect sizes were computed for the age effect in each individual model. The change in effect size of the age covariate between the single regressor age model and the age-plus-covariate models was computed for each covariate.

### 4.8 Effect Size and Power Analyses

Effect sizes are computed using Cohen’s *F*^2^ metric, which compares the *R*^2^ of two models to establish the contribution of a particular regressor to the variance explained by the model [Cohen, 1988, Selya et al., 2012]. Cohen’s *F*^2^ can be interpreted as the proportion of residual variance explained by the addition of the new regressor.

The effect size is defined as:

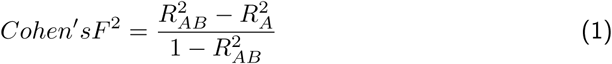

In which B refers to the regressor of interest and A refers to the set of all other regressors in the model. The numerator is then the proportion of additional variance explained by addition of the regressor of interest B. This is normalised by the denominator, which quantifies the total residual variance left unexplained by the full model [Selya et al., 2012]. A full spectrum of effect sizes can be computed from the GLM-Spectrum model [Quinn et al., 2024].

Bootstrapped confidence intervals for effect size estimates were computed by building a distribution of effect sizes using 1000 iterations of resampling with replacement. 95% confidence intervals were computed as percentiles across the bootstrapped distribution of effect size estimates. Confidence intervals for power and sample size calculations are computed using the bootstrapped Cohen’s *F*^2^ metrics.

Power analysis for regressors in a general linear model was computed using in-house Python code validated against the pwr R package [Champely, 2020, Cohen, 1988]. Five variables were defined, of which any four can be used to compute the remaining fifth variable. For a GLM, these variables are the number of observations in the data, the number of regressors in the model, the estimated effect size, the alpha level (type 1 error), and the power (1 - type 2 error). We computed the future sample size estimates in Tables 1 and 2 using this approach. The number of observations in the dataset were computed from the number of regressors in the model (2 or 3), the Cohen’s *F*^2^ estimate, an alpha level of 0.05, and a power of 80%.

## Data & Code Availability

Raw sensor-level MEG recordings are from the Cam-CAN Data Repository: https://opendata.mrc-cbu.cam.ac.uk/projects/camcan/ (Eyes Closed Resting State data) and the MEG-UK Database https://meguk.ac.uk/database/ (Eyes closed resting data from Oxford, Cambridge & Nottingham). Preprocessed and analysed data to reproduce group-level analyses are archived on the Open Science Framework : https://osf.io/yspdg/. Code to run all processing, data analysis and visualisations are available on github: https://github.com/OHBA-analysis/Quinn2025-RobustReplicableAge.

The analyses in this paper were carried out in Python 3.11 with core dependencies as numpy [Harris et al., 2020], scipy [Virtanen et al., 2020] and Matplotlib [Hunter, 2007]. MNE python [Gramfort, 2013] was used for EEG/MEG data processing with OSL batch processing tools [Quinn et al., 2022, van Es et al., 2025]. The spectrum analyses further depend on the Spectrum Analysis in Linear Systems toolbox [Quinn and Hymers, 2020] and glmtools^1^.

## Funding

This research was supported by the National Institute for Health Research (NIHR) Oxford Health Biomedical Research Centre. The Wellcome Centre for Integrative Neuroimaging is supported by core funding from the Wellcome Trust (203139/Z/16/Z). CG is supported by the Wellcome Trust (215573/Z/19/Z). OK is supported by the Marie Sklodowska-Curie Innovative Training Network “European School of Network Neuroscience (euSNN)” (860563). MWJvE is supported by the Wellcome Trust (106183/Z/14/Z, 215573/Z/19/Z), the New Therapeutics in Alzheimer’s Diseases (NTAD) supported by the MRC and the Dementia Platform UK (RG94383/RG89702). ACN is supported by the Wellcome Trust (104571/Z/14/Z) and James S. McDonnell Foundation (220020448). MWW is supported by the Wellcome Trust (106183/Z/14/Z, 215573/Z/19/Z), the New Therapeutics in Alzheimer’s Diseases (NTAD) study supported by UK MRC, the Dementia Platform UK (RG94383/RG89702) and the NIHR Oxford Health Biomedical Research Centre (NIHR203316). The views expressed are those of the author(s) and not necessarily those of the NIHR or the Department of Health and Social Care.

For the purpose of open access, the author has applied a CC-BY public copyright licence to any Author Accepted Manuscript version arising from this submission.

The computations described in this paper were performed using the University of Birmingham’s BlueBEAR HPC service, which provides a High Performance Computing service to the University’s research community. See http://www.birmingham.ac.uk/bear for more details.

## Author Contributions

A.J.Q.: Conceptualisation, Methodology, Software, Formal analysis, Data Curation, Project administration, Writing—Original Draft, Writing—Review & Editing, and Visualisation.

C.G.: Conceptualisation, Methodology, Software, Formal analysis, Data Curation, and Writing—Review & Editing. J.P.: Conceptualisation, Formal analysis and Writing—Review & Editing O.K.: Conceptualisation, Writing—Original Draft & Writing—Review & Editing M.v.E.: Conceptualisation, Writing—Review & Editing. A.C.N.: Conceptualisation, Writing—Review & Editing, Supervision. M.W.W.: Conceptualisation, Methodology, Writing—Original Draft, Writing—Review & Editing, and Supervision.

## A Supplementary Material

### A.1 Literature review of alpha effects

Part of our motivation for this method is that variability in the methodological choices made in different publications makes it difficult to aggregate varying results across the literature. For example, though decrease in alpha peak frequency with increasing age is reported highly consistently, the effect of age on alpha power is much more variable. Table 3 shows a sample of publications over the last 20 years that report a change in alpha power with age (note that this is intended to be a representative rather than an exhaustive list). Over half of publications (9/15) report a decrease in alpha power whilst the remaining publications report an increase (2/15), both increases and decreases (1/15), a decrease but only without correcting for aperiodic slope (1/15), a quadratic effect (1/15) and no-effect (1/15). Stier et al. [2023] suggest that the choice of analysis space (sensor space or source reconstruction) might drive these differences.

**Table 3.** A selection of publications from the last 20 years reporting age-dependent differences in alpha power. A variety of methodological approaches are used by the papers. Note that this is not intended to be a comprehensive list of all publications in this time, but is a representative sample of the variability in methods and results found in the literature.

| Publication | Method / Frequency / Location | Change with age | Figure |
| --- | --- | --- | --- |
| Babiloni et al. [2006] | Method: MEG source space<br>Alpha: 8–10.5 Hz; 10.5–13 Hz<br>Peak: Parietal, occipital, temporal, limbic areas | Decrease | Figures 1, 2 & 3 |
| Lodder and van Putten [2011] | Method: EEG sensor space<br>Alpha: Individual peak (3–18Hz)<br>Peak: Individual peak | Decrease | Table 2 2 |
| Vysata et al. [2012] | Method: EEG sensor space<br>Alpha: 7–13 Hz<br>Peak: Whole head | Decrease | Figure 2 |
| Gómez et al. [2013] | Method: MEG sensor space<br>Alpha: 8–13 Hz<br>Peak: Parieto-temporal areas | Quadratic | Figure 3 |
| Scally et al. [2018] | Method: EEG sensor space<br>Alpha: Individual peak (6–14 Hz) and upper alpha (10–12Hz)<br>Peak: All channels averaged | No change (individual)<br>Decrease (upper alpha) | Figure 5 |
| Hoshi and Shigihara [2020] | Method: MEG source space<br>Alpha: 8–12 Hz<br>Peak: Temporal pole; Visual cortex V1–V4 | Increase (TP)<br>Decrease (Visual) | Table 3 |
| Donoghue et al. [2020] | Method: EEG sensor space<br>Alpha: Individual peak (7–14 Hz)<br>Peak: Sensor Oz | Decrease | Figure 5 |
| Thuwal et al. [2021] | Method: MEG sensor space<br>Alpha: Individual peak (8–12 Hz)<br>Peak: Occipito-parietal sensors | Decrease | Figures 6 |
| Rempe et al. [2023] | Method: MEG source space<br>Alpha: 8–12 Hz<br>Peak: Bilateral inferior temporal gyrus | Increase | Figures 2 & 3 |
| Stier et al. [2023] | Method: MEG source space<br>Alpha: 8–12 Hz<br>Peak: Temporal regions | Increase | Figure 2 |
| Merkin et al. [2023] | Method: EEG sensor space<br>Alpha: Individual peak (after 1/f correction)<br>Peak: All channels | No change | Figure 3 |
| Tröndle et al. [2023] | Method: EEG sensor space<br>Alpha: Individual peak $\pm 2$Hz (8–13 Hz)<br>Peak: Parietal and occipital | Decrease | Figures 5 & 7 |
| Park et al. [2024] | Method: EEG source space<br>Alpha: Individual peak (6 to 14 Hz; after 1/f correction)<br>Peak: Sensorimotor & parietal regions | Decrease | Figures 2 & 3 |
| Ustinin et al. [2025] | Method: MEG sensor space<br>Alpha: 8–13Hz<br>Peak: Whole head | Decrease | Figures 5 & 6 |
| Medrano et al. [2025] | Method: EEG sensor space<br>Alpha: Individual peak (8–12Hz; after 1/f correction)<br>Peak: Channel Oz | Decrease | Figures 8 |

Critically, it is difficult to reconcile these findings with the information reported in the publications. For example, even within the 9 publications that report a decrease, there is little correspondence in the spatial location of the effect. We argue that focused approaches cannot resolve this issue alone as targeted analyses are more specific to each dataset and less generalisable.

In the specific case of the mixed literature on change in alpha power with age, our results show that both effects are present and separated in frequency. It is feasible that the different methodological choices and datasets used by each study in our survey means that one or other of these two effects were emphasised. As a result, the literature may not be mixed in scientific terms, but that a rich pattern of results is obscured by methodological variability.

Importantly, the differences in alpha power may be explainable with the change in alpha frequency. Though this was not explicitly modelled by our analysis, it can be visualised by quantifying the alpha peak in the model fitted spectra. A strong reduction in alpha peak frequency is qualitatively visible in the model projected power spectrum (Figure 9A). Whilst no single parameter directly fits this feature in the GLM-Spectrum, this pattern is captured by the profile of linear parameters across channels and frequencies. This pattern contains both and increase and decrease in magnitude with age, separated by frequency (Figure 9B). The decrease in alpha peak frequency can be recovered by extrema detection carried out on the model projected spectra (Figure 9A) and shows a prominent decrease in alpha power and alpha frequency across the course of ageing. Critically, by the full profile of parameter estimates the shape of overall spectrum can represent non-linear changes features that are derived from the full spectrum (Figure 9C & D).

**Figure 9.**
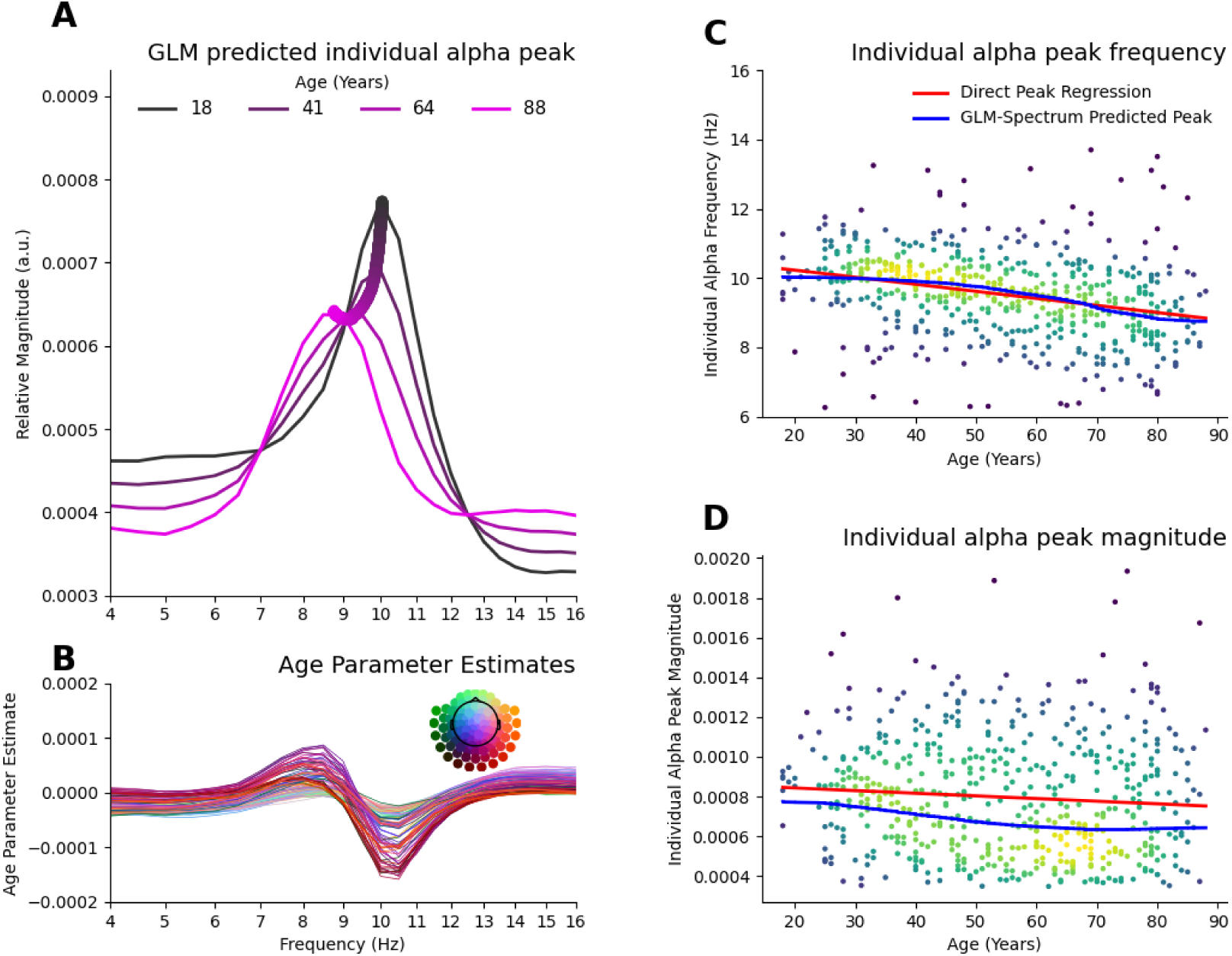
Model projected age effect retains non-linear properties of decrease in alpha peak frequency and magnitude. **A)** Model predicted spectra (averaged across all sensors) for the oldest and youngest participants in Cam-CAN with the alpha peak for intermediate ages overlaid in the thick line. **B)** The parameter estimates of age that combine to describe the peak frequency shift seen in A). **C)** Scatter plot of individual alpha peak frequencies against age. A linear regression line fitted directly to the individual alpha peaks is shown in red and the GLM-Spectrum derived alpha peaks from the model projected spectra is shown in blue. **D)** As C for individual alpha peak magnitude.

### A.2 Source space age effect

The sensorspace effect shown in Figure 1 is replicated in a source projected analysis using an LCMV beamformer with voxel data organised into parcels are orthogonalised. The average profile of the age effect in source space is highly consistent with the sensorspace result.

**Figure 10.**
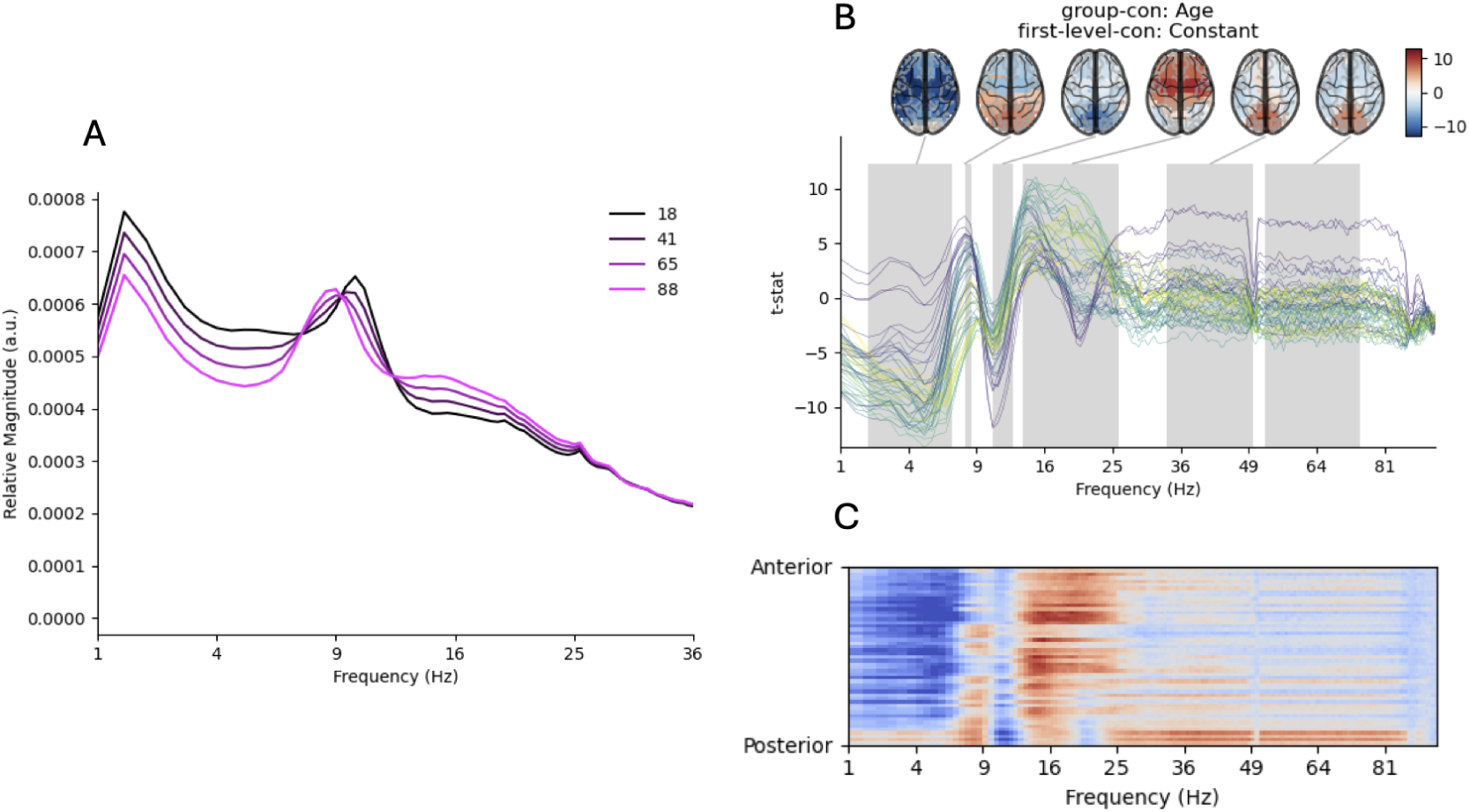
Effect of age on the relative magnitude spectrum across space and frequency in LCMV source reconstructed and parcellated data. **A)** Model-predicted spectra (averaged across all sensors) for 4 equally spaced ages across the participant age range. **B)** Spectrum of t-values quantifying the age effect across space and frequency. Source topographies are shown in the frequency bands with significant effects at sensorspace. The permutation statistics are not repeated at source space. **C)** A 2D frequency-by-space map of all statistically significant effects. Sensor-Frequency combinations that do not reach statistical significance have a faded colour scale. Blue regions indicate decreasing spectral power with age and red regions indicate increases.

### A.3 Effect of ICA denoising on the age effect

The age results may be contaminated in some way by residual cardiac or ocular artefacts that are not removed during preprocessing. Though ICA denoising was applied, it is possible that some artefactual components were not identified and removed from the dataset. The results at low frequencies and in the gamma range are most likely to be directly impacted by this contamination.

To explore the impact this has on our analysis, we reran the core GLM effect of age on the data with no ICA artefact rejection at all, allowing all eye movements and heart rate components to remain in the data (Figure 11).

**Figure 11.**
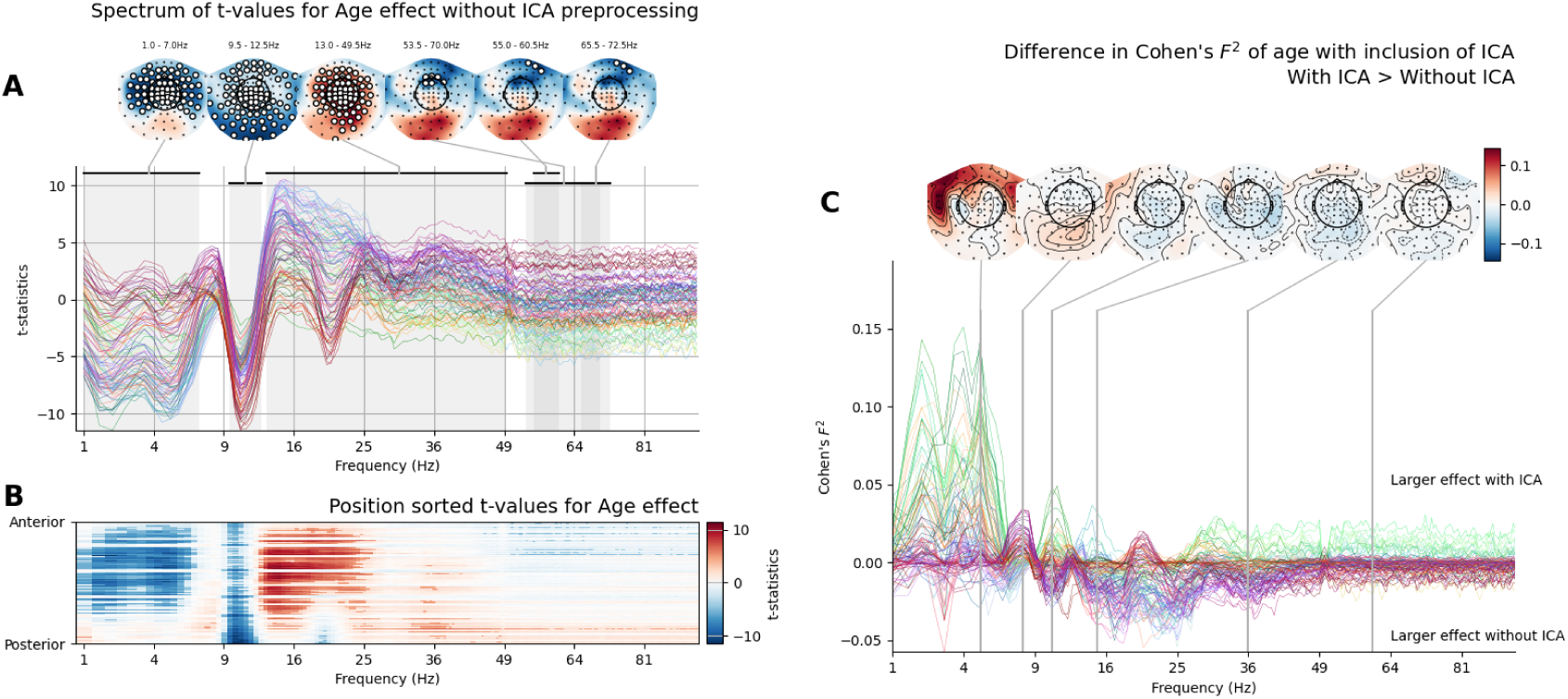
Effect of age with and without ICA denoising is largely consistent. **A)** Spectrum of t-values quantifying the age effect across space and frequency. Non-parametric permutations with maximum statistics to control for multiple comparisons across sensors and frequency bins. While this permutation testing was not cluster-based, contiguous clusters of significant sensors were computed post-hoc for visualisation The largest 6 spatially and spectrally contiguous areas of statistically significant effects are highlighted in frequency by black bands at the top of the spectrum and highlighted in space by sensors marked with a white circle in the adjoining topography. **B)** A 2D frequency-by-space map of all statistically significant effects. Sensor-Frequency combinations that do not reach statistical significance have a faded colour scale. Blue regions indicate decreasing spectral power with age and red regions indicate increases. **C)** Spectrum of the difference in estimated effect size of age ind ata that has been cleaned with ICA and data that has not be cleaned with ICA. Positive values indicate that the effect was larger with ICA denoising and negative values indicated that the effect was smaller with ICA.

This no-ICA analysis has three differences to the original in the publication: The low-frequency decrease with age is stronger and more widespread in the analysis that removes ocular artefacts with ICA. A large negative effect is visible in both analyses, though without ICA several frontal and temporal sensors no longer show significant effects. Similarly, the effect size of the low frequency effect of age is substantially larger when ICA denoising is applied.

The low alpha effect was strongly reduced in the analyses that do not remove artefacts with ICA. A large central-occipital group of sensors show an effect between 7 and 8.5Hz in the ICA analysis, but this is reduced to a single sensor at 8Hz when ICA is not computed. At high frequencies, the age effect in frontal sensors is larger with ICA and the age effect in posterior sensors is larger without ICA, though the position and frequencies of significant effects are largely unchanged. The remaining effects in the high alpha and beta ranges are unchanged by application of ICA.

Overall, ICA either improves the estimation of age effects (low-frequency, low-alpha, high-gamma) or has negligible effect (high-alpha, beta). Only the posterior low-gamma effect is reduced by ICA. Together, we take this as evidence that our core results are robust to interference by eye movements and that the ICA denoising is working effectively to reduce noise in the analysis.

### A.4 Details of covariate analyses

The GLM-Spectrum effects of the investigated covariate effects are summarised in Figure 7 and shown here in full detail. Figures 12, 13 and 14 shows the results for models fitted with an intercept term and the z-transformed covariate. The full GLM-Spectrum is shown for the parameter estimates (Figure 12), t-statistics (Figure 13) and Cohen’s *F*^2^ effect sizes (Figure 14). The results from Figure 14 are summarised in Figure 7A in the main text.

**Figure 12.**
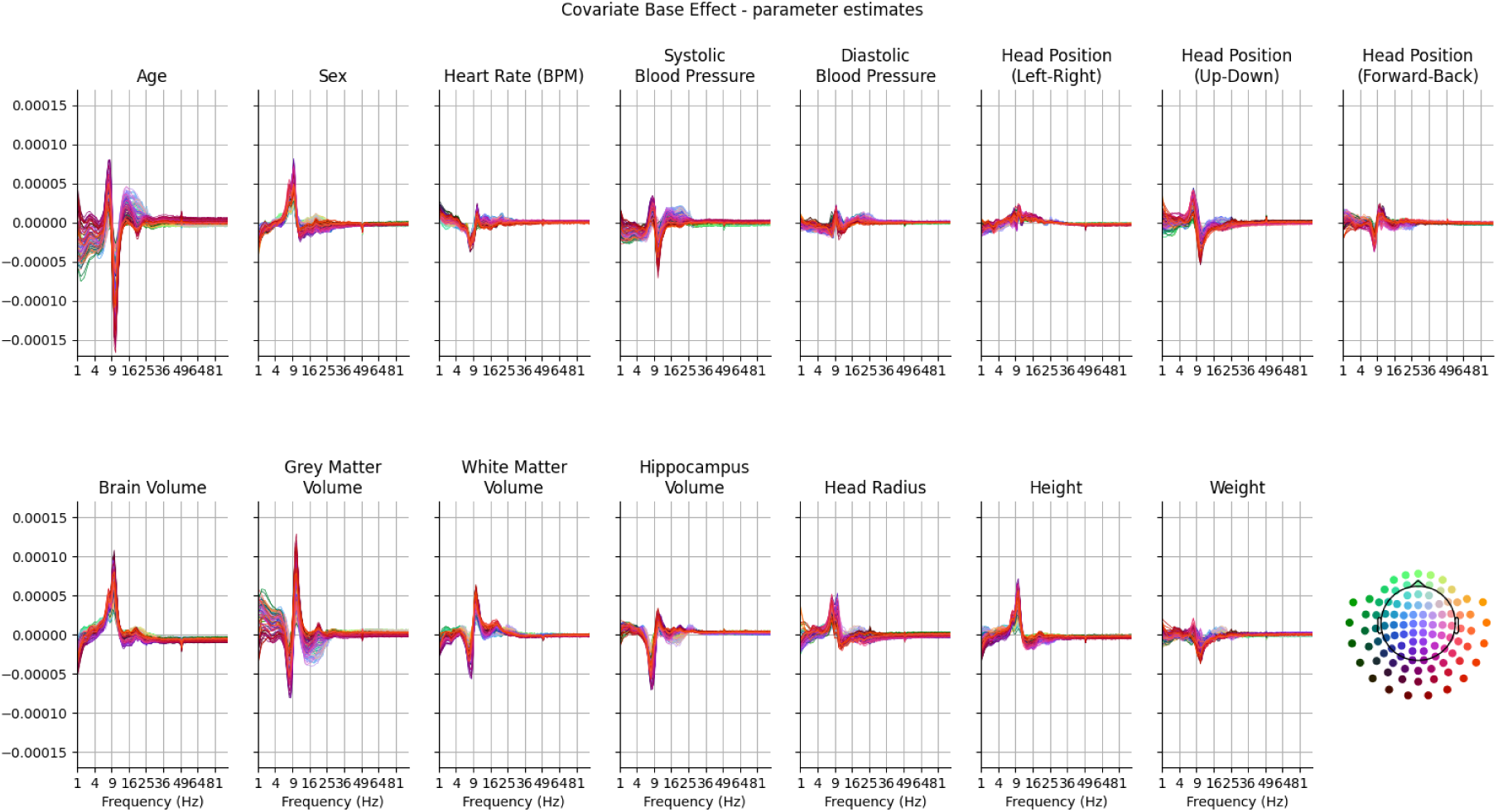
GLM spectrum of parameter estimates for all covariates.

**Figure 13.**
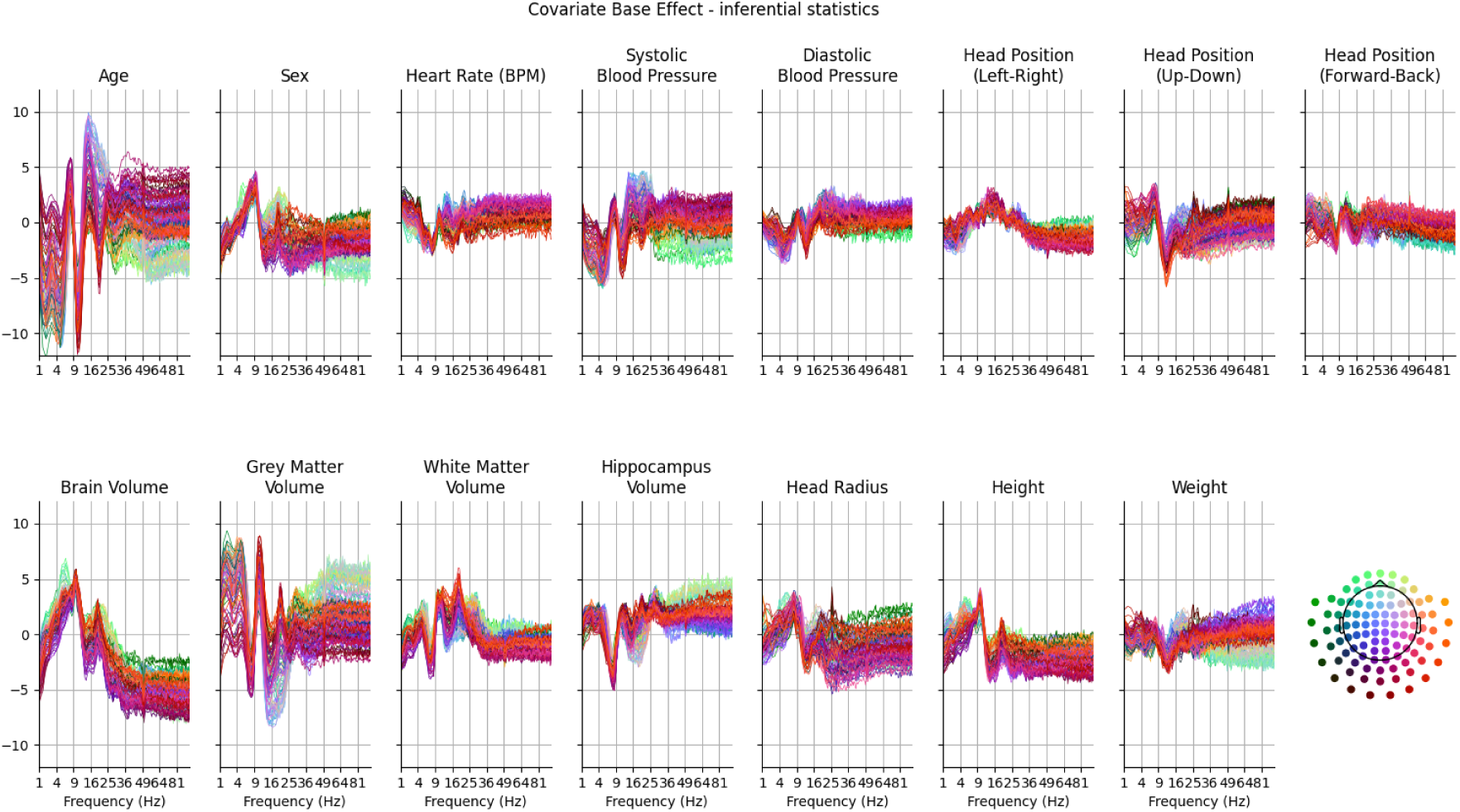
GLM spectrum of t-statistics for all covariates.

**Figure 14.**
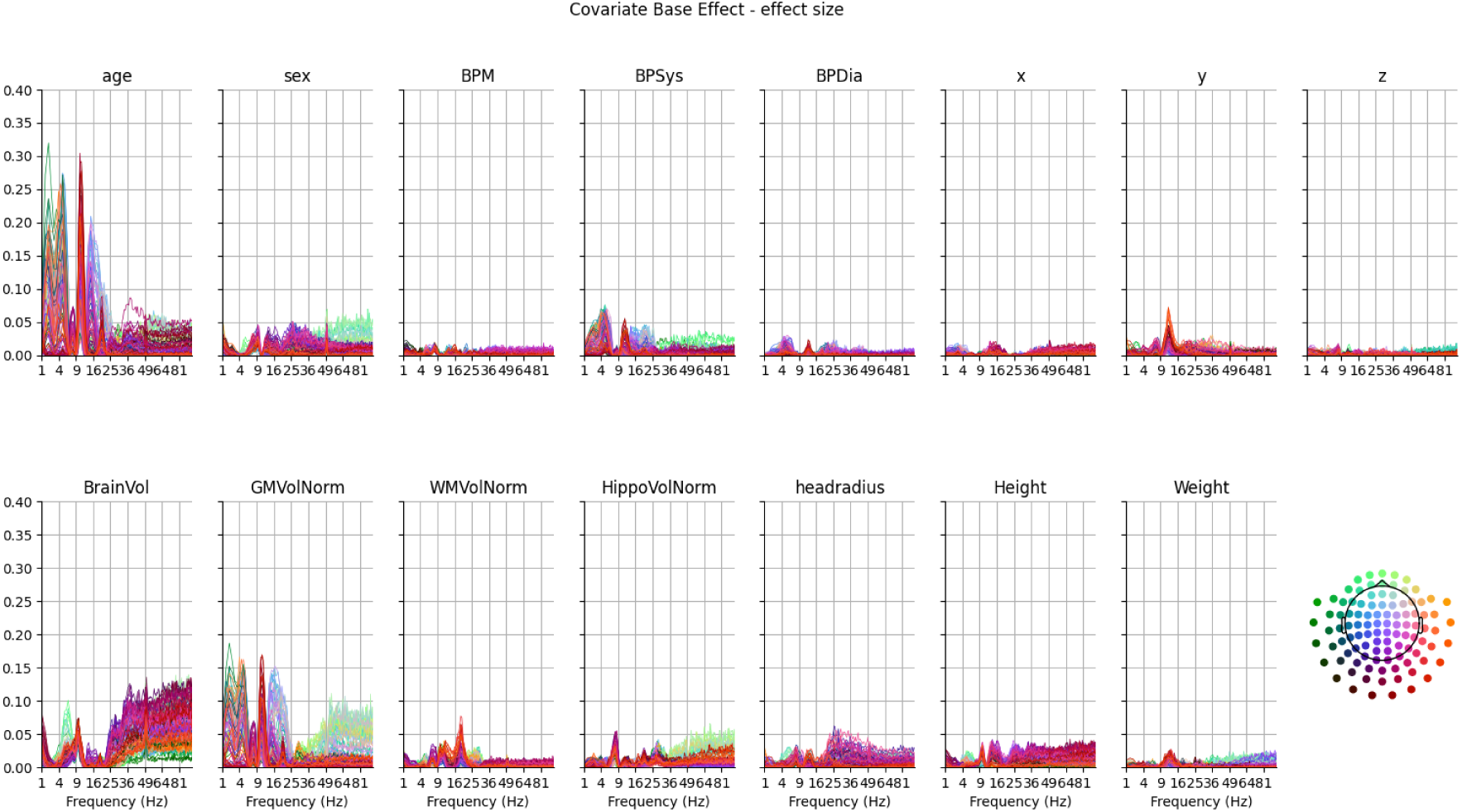
GLM spectrum of effect sizes for all covariates.

Figure 15 shows how the Cohen’s *F*^2^ estimates for age change between two models. The first model contains an intercept and the z-transformed ages as regressors and the second model contains the z-transformed values of an additional covariate. Cohen’s *F*^2^ for the age regressor is computed from both models and Figure 15 shows the difference. Age itself is excluded from this analysis. This result is summarised in Figure 7B in the main text.

**Figure 15.**
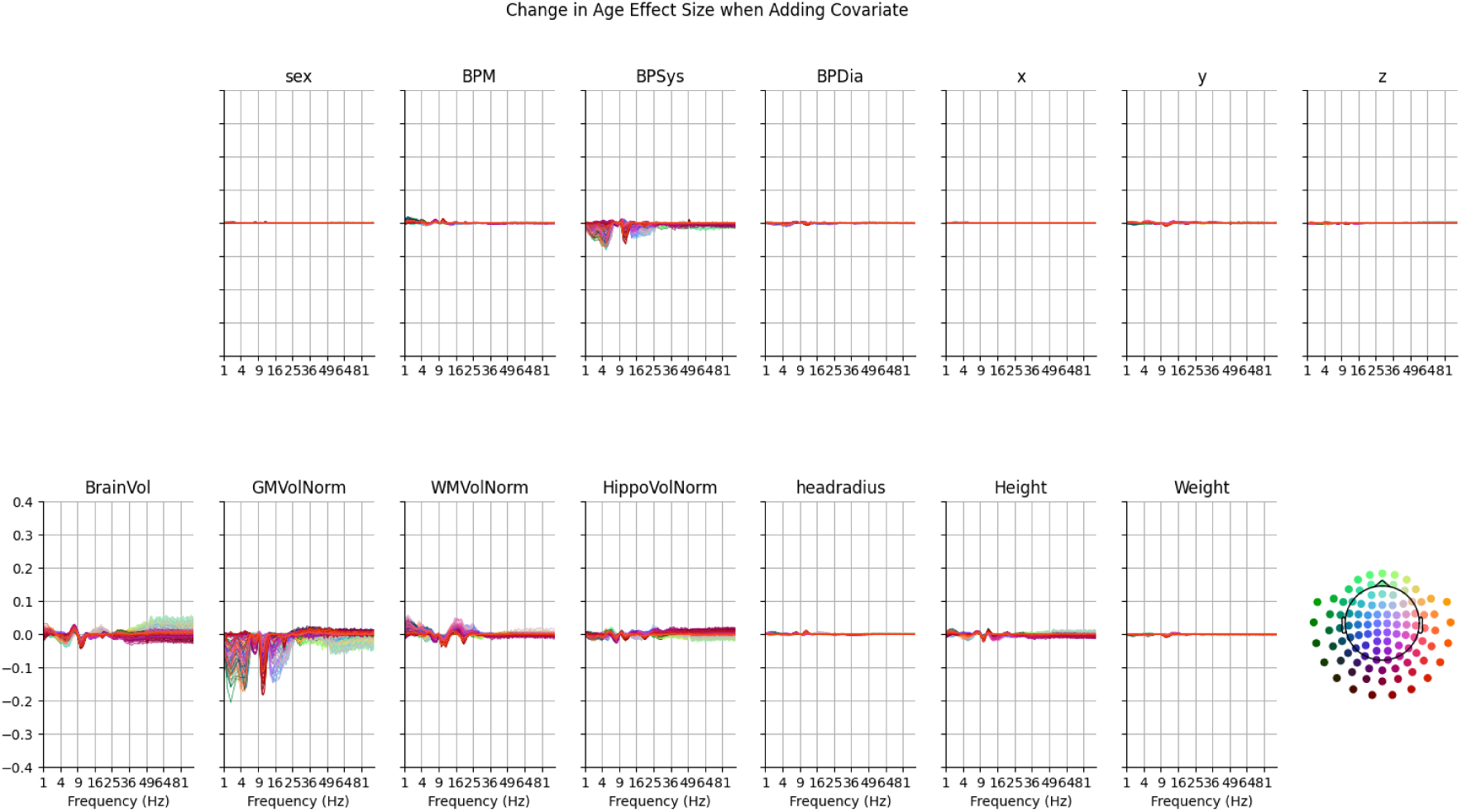
GLM spectrum of change in estimates age effect size when including covariate in the model. Age itself is excluded from this figure.

### A.5 Robustness to head position correction methods

The head position of participants within the MEG sensor dewar is an important consideration that has the potential to change the signal-to-noise level of each individual data recording. There is a significant difference in head position as a function of age in the CamCAN dataset in Y (front-back) direction indicating that older participants are seated further forward in the dewar than younger participants.

**Figure 16.**
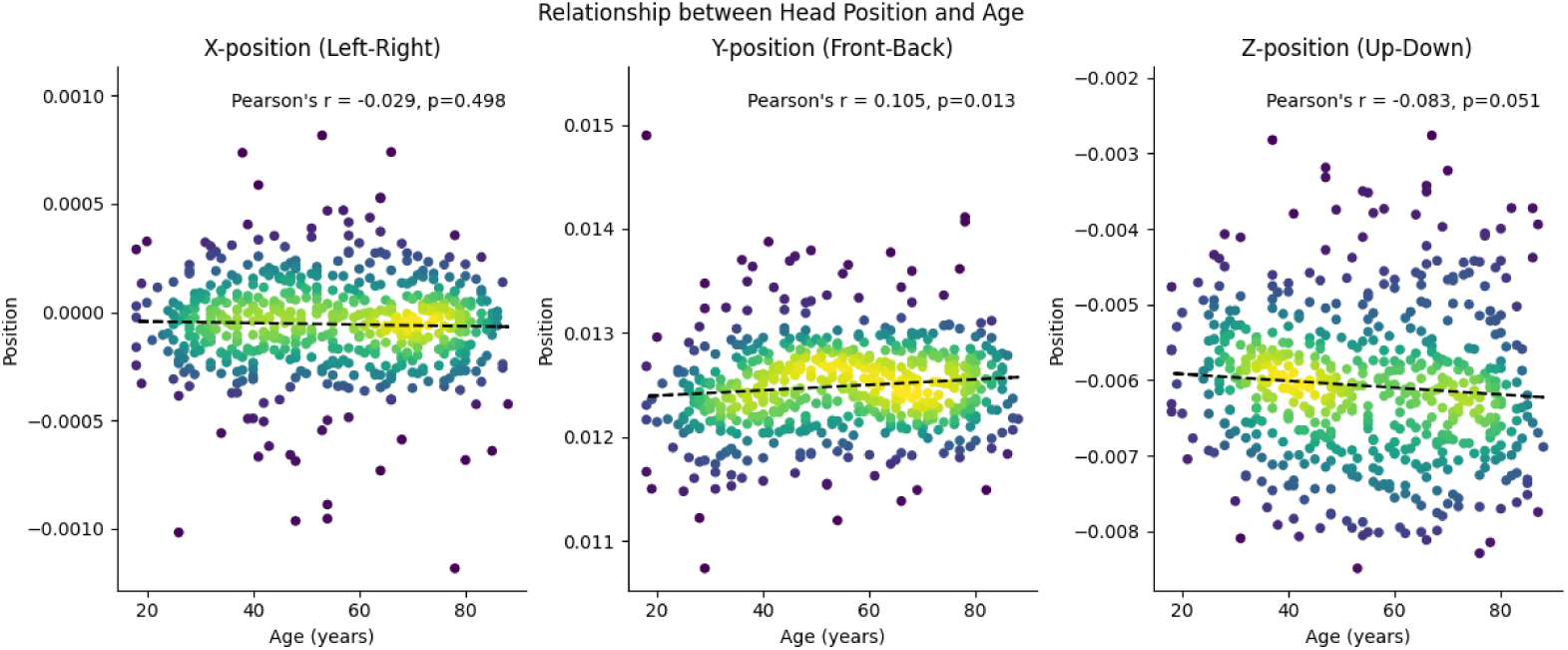
Correlations between head position and age for the CamCAN dataset.

Both maxfilter processing and transformation to relative power are thought to at least partially compensate for differences in head position. The change in the GLM spectrum with respect to head position correction is discussed in Section 2.4. The results in Figure 17 show the full GLM-Spectra for the age effect computed for each combination of two sensor normalisation conditions - no normalisation (absolute magnitude) and z-transformed time series (relative magnitude) - and three SSS based head position correction conditions - no correction, within recording head position correction (*−movecomp*) and within and between recording head position correction (*−movecomp* and *−trans*). Results from the analysis of the relative magnitude with both within and between recording head position correction are shown in Figure 1 of the main text. Results from the analysis of the absolute magnitude with both within and between recording head position correction are shown in Figure 5 of the main text The SSS head position correction methods have little effect on the estimated group age effect whereas the sensor normalisation change the overall pattern of results as discussed in Section 2.4.

**Figure 17.**
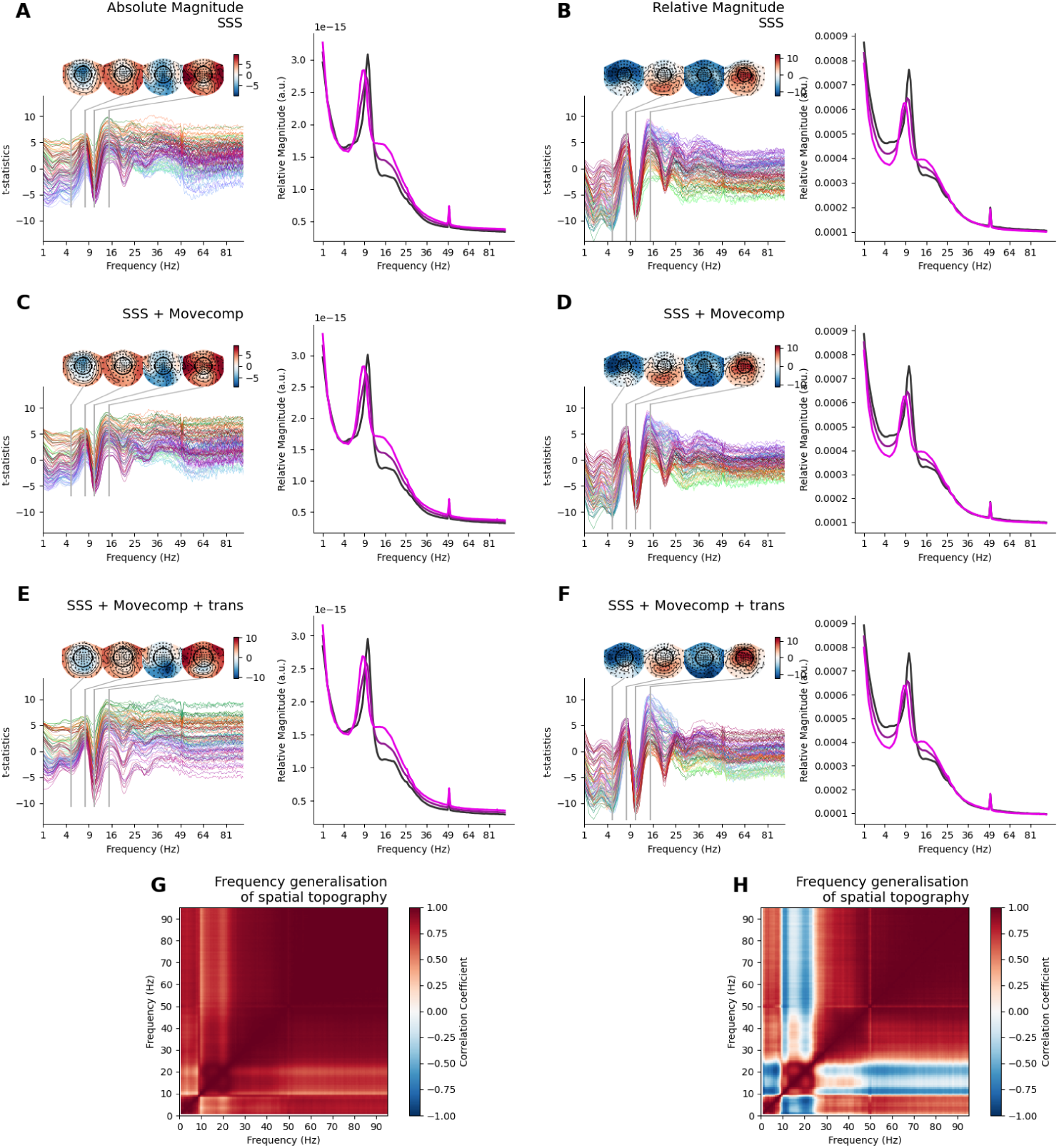
GLM-Spectrum results showing the effect of age estimated for two different sensor normalisations and three different SSS head position correction types. **A)** The age effect for the absolute magnitude with SSS applied without head position correction. **B)** The age effect for the relative magnitude with SSS applied without head position correction. **C)** The age effect for the absolute magnitude with SSS applied with head position correction applied within each dataset (*−movecomp* in maxfilter software). **D)** The age effect for the relative magnitude with SSS applied with head position correction applied within each dataset (*−movecomp* in maxfilter software). **E)** The age effect for the absolute magnitude with SSS applied with head position correction applied within each dataset (*−movecomp* in maxfilter software) and head position alignment to a reference datafile (*−trans* in maxfilter software). **F)** The age effect for the relative magnitude with SSS applied with head position correction applied within each dataset (*−movecomp* in maxfilter software) and head position alignment to a reference datafile (*−trans* in maxfilter software). **G)** The spectral generalisation of the topography of the absolute magnitude age effect. **H)** The spectral generalisation of the topography of the relative magnitude age effect.

The impact of the sensor normalisation is a change in the spectral specificity of spatial patterns in the age effect. The largest component of the absolute magnitude age effect is a consistent spatial pattern covering the whole frequency range, leading to high positive correlations in the spatial maps of the age effect across all frequencies (Figure 17G). In contrast, spatial topography of the relative magnitude age effect varies strongly across frequency (Figure 17H)

### A.6 Multicollinearity between Age and Grey Matter Volume

There is a strong correlation between age and Global Grey Matter Volume (Pearson’s r=-0.75). The shared variance arising from this collinearity adds nuance to the interpretation of the results, which we explore in more detail in this section.

Firstly, the sum-square residuals for the group-level model fit including both Age and GGMV are shown as a function of frequency in Figure 18. We see that the residuals broadly follow the overall distribution of variance in the data, peaking at low frequencies and in the alpha range.

**Figure 18.**
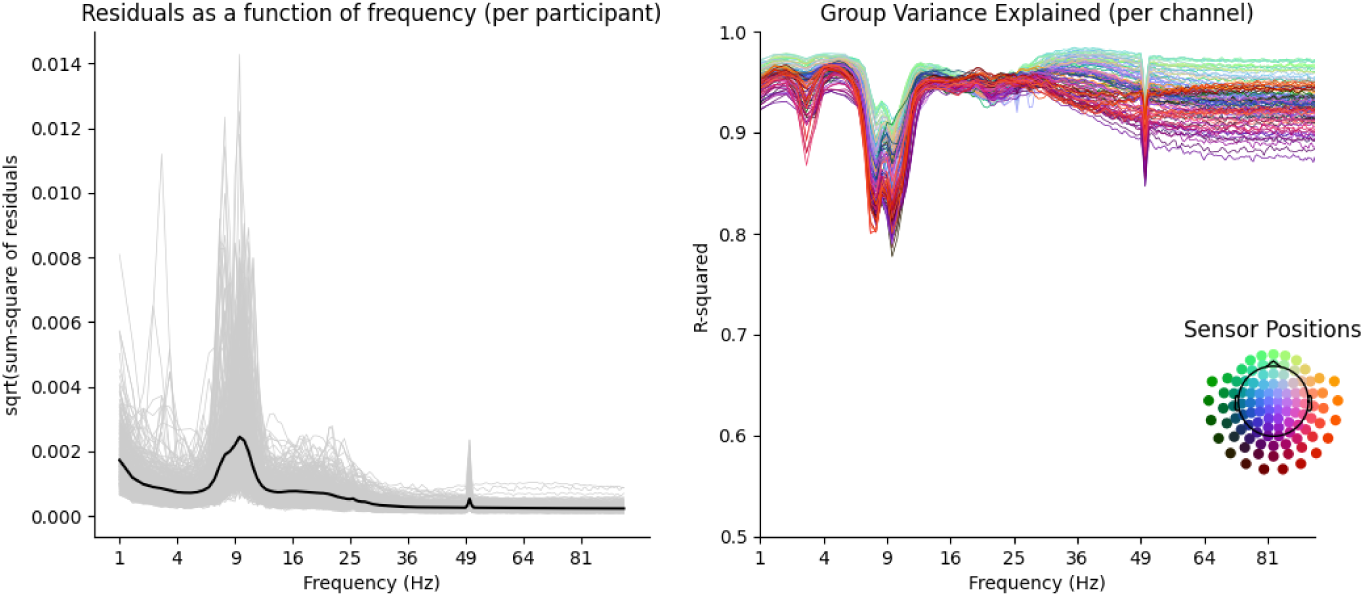
Model residuals and coefficient of determination as a function of frequency for each channel show that the group model performs well. Left) The sum-square residuals of the first level models with each individual data recording (grey lines) and the overall group average (black line). Right) The *R*^2^ coefficient of determination for the group-level model showing that the model is able to describe a high amount of variability across both channels and frequencies.

These frequency ranges are where the strongest signal is visible, but also the highest variability between participants. We would expect that the group model would not perform so well in the points of greatest variability. Importantly, though the residuals are relatively high in the alpha this is still in the context of a very well-performing model with R2 values of around 80%.

Secondly, the correlation between age and GGMV is not inherently problematic for the GLM, but it does add complexity and nuance to the interpretation of the results. Some additional model validation statistics are shown in Figure 19.

**Figure 19.**
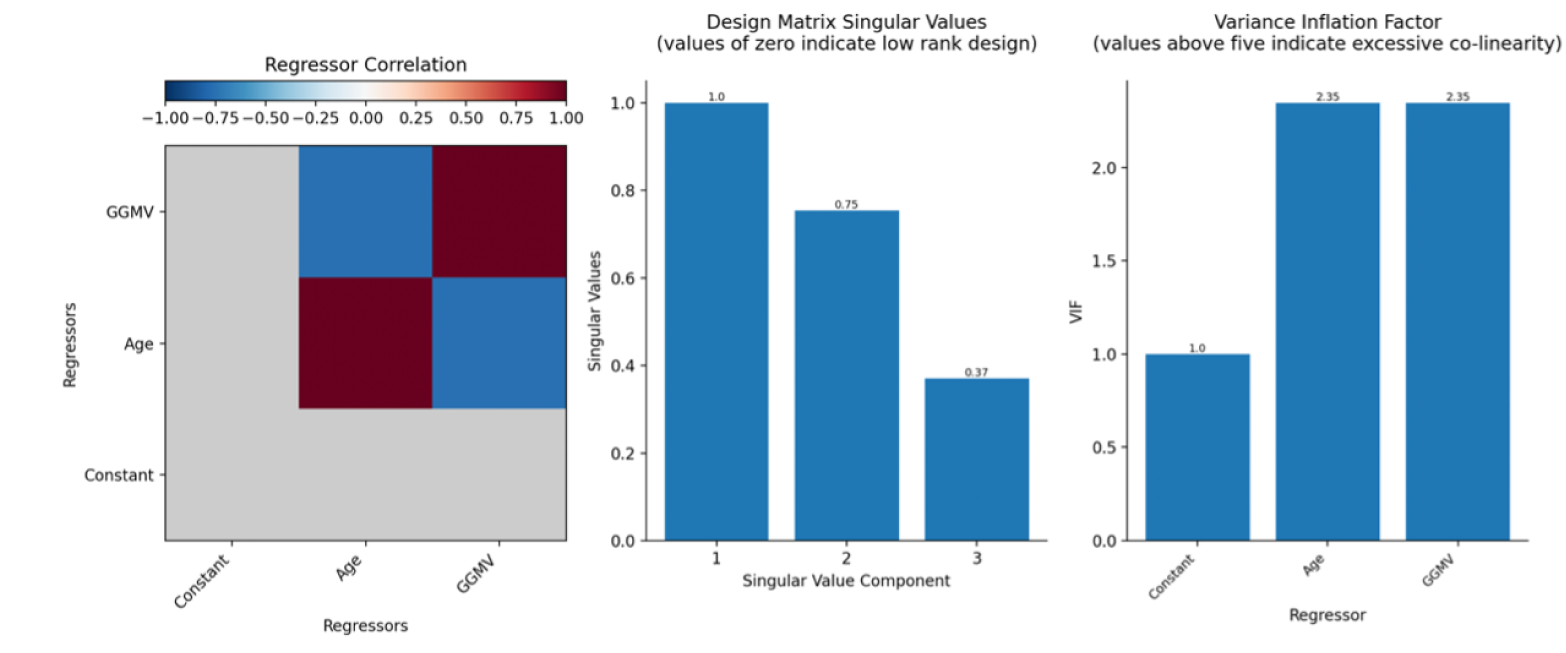
Model validation statistics and diagnostics for the group-level GLM containing an intercept, a z-transformed age regressor and a z-transformed Global Grey Matter Volume regressor. Left) The correlation matrix between the three regressors shows a large negative correlation between age and GGMV. Centre) The singular values of the design matrix stay well above zero for all components, indicating that the collinearity between regressors does not cause the model fit to critically fail. Right) The variance inflation factors for all three regressors shows peak values of 2.35 which indicates inflated variance (VIF ¿ 1). Specifically, this corresponds to an increase in standard error by a factor of 1.533.

The singular value spectrum of the design matrix indicates whether a design is low-rank, the smallest singular value in this case in 0.37 which indicates that there isn’t a rank deficiency which would prevent us from estimating the model.

Though we can estimate the model, correlated regressors can reduce its efficiency. The variance inflation factors for the joint AGE-GGMV model are above 1 for both parametric regressors, indicating that the standard errors of their estimates are inflated. The VIF of 2.35 indicates that the standard errors of this joint model are around sqrt(2.35) = 1.533 times greater than they would be in a separate or uncorrelated model. Though there is no hard rule for this, the literature generally suggests that a VIF above 5 (or sometimes 10) indicates severe multicollinearity.

Including additional regressors in the model can change the age estimate by ‘partialling’ out the variance that can be attributed to the other variables, and by inflating standard errors. In this specific case of GGMV, the partialled estimates are reduced heterogeneously across space and frequency, and the amount of inflation is at a tolerable level. Overall, the regression is able to separate the unique effects of age and GGMV, at the cost of this inflation in the associated standard errors

Finally, the partialled effects in the GLM can already be interpreted as the “change in Y for a 1-unit increase in X whilst all other predictors are held constant”. In our case, this is equivalent to asking “Is there an effect of age on the MEG spectrum in ‘a group where GGMV does not change with age”. The results show that it is not feasible to interpret changes in low-alpha separately from change in GGMV, but that a substantial component of the low-frequency, high alpha and beta band changes are distinct from global structural change.

### A.7 Analysis Family

**Table 4.** Summary of regression analyses performed in this article organised by figure.

| Figure | Field | Specification |
| --- | --- | --- |
| 1, 2, 4, 9, 18, 19 | Analysis<br>Dimensions<br>Model<br>Permutations | Linear effect of age (Cam-CAN)<br>Sensors × frequency<br>$RelativePower = Intercept + ztrans(Age)$<br>2500 |
| 3 | Analysis<br>Dimensions<br>Model<br>Permutations | Quadratic effect of age (Cam-CAN)<br>Sensors × frequency<br>$RelativePower = Intercept + ztrans(Age) + ztrans(Age)^2$<br>2500 |
| 5 | Analysis<br>Dimensions<br>Model<br>Permutations | Linear effect of age (Cam-CAN & MEG-UK)<br>Sensors × frequency<br>$RelativePower = Intercept + ztrans(Age)$<br>2500 |
| 6, 17 | Analysis<br>Dimensions<br>Model<br>Permutations | Linear effect of age (Cam-CAN)<br>Sensors × frequency<br>$AbsolutePower = Intercept + ztrans(Age)$<br>2500 |
| 7, 12, 13, 14 | Analysis<br>Dimensions<br>Model<br>Permutations | Linear effect of covariates (Cam-CAN)<br>Sensors × frequency<br>$RelativePower = Intercept + ztrans(Covariate)$<br>2500 |
| 7, 12, 13, 14 | Analysis<br>Dimensions<br>Model<br>Permutations | Linear effect of age with covariate included (Cam-CAN)<br>Sensors × frequency<br>$RelativePower = Intercept + ztrans(Age) + ztrans(Covariate)$<br>2500 |
| 8, 9 | Analysis<br>Dimensions<br>Model<br>Permutations | Linear effect of GGMV with age (Cam-CAN)<br>Sensors × frequency<br>$RelativePower = Intercept + ztrans(Age) + ztrans(GGMV)$<br>2500 |
| 10 | Analysis<br>Dimensions<br>Model<br>Permutations | Linear effect of age (Cam-CAN)<br>Source parcels × frequency<br>$RelativePower = Intercept + ztrans(Age)$<br>2500 |
| 11 | Analysis<br>Dimensions<br>Model<br>Permutations | Linear effect of age without ICA (Cam-CAN)<br>Sensors × frequency<br>$RelativePower = Intercept + ztrans(Age)$<br>2500 |

## Footnotes

1 https://pypi.org/project/glmtools/

